# Action perception recruits the cerebellum and is impaired in spinocerebellar ataxia patients

**DOI:** 10.1101/480293

**Authors:** Abdel R. Abdelgabar, Judith Suttrup, Robin Broersen, Ritu Bhandari, Samuel Picard, Christian Keysers, Chris I. De Zeeuw, Valeria Gazzola

**Affiliations:** Social Brain Lab & Cerebellar Coordination and Cognition Group, Netherlands Institute for Neuroscience, A research Institute of the Royal Netherlands Academy of Arts and Sciences. Amsterdam, The Netherlands; Department of Neuroscience, University Medical Centre Groningen, University of Groningen. Groningen, The Netherlands; Brain & Cognition, Department of Psychology, University of Amsterdam. Amsterdam, The Netherlands; Department of Neuroscience, Erasmus Medical Center, Rotterdam, The Netherlands

**Author notes:** Equal contribution (Abdel R. Abdelgabar, Judith Suttrup, Robin Broersen, Ritu Bhandari). Equal contribution (Christian Keysers, Chris I. De Zeeuw, Valeria Gazzola). **Corresponding author:** Valeria Gazzola Social Brain Lab, Netherlands Institute for Neuroscience Meibergdreef 47, 1105 BA Amsterdam, The Netherlands.

## Abstract

Our cerebellum has been proposed to generate prediction signals that may help us plan and execute our motor programs. However, to what extent our cerebellum is also actively involved in perceiving the action of others remains to be elucidated. Using fMRI, we show here that observing goal-directed hand actions of others bilaterally recruits cerebellar Lobules VI, VIIb and VIIIa. Moreover, whereas healthy subjects (n=31) were found to be able to discriminate subtle differences in the kinematics of observed limb movements of others, patients suffering from spinocerebellar ataxia type 6 (SCA6; n=21) were severely impaired in performing such tasks. Our data suggest that the human cerebellum is actively involved in perceiving the kinematics of the hand actions of others and that SCA6 patients’ deficits include a difficulty in perceiving the actions of other individuals. This finding alerts us to the fact that cerebellar disorders can alter social cognition. Given that impairments in social cognition have been reported to be one of the most debilitating consequences of neurological disorders, this finding may be relevant to improving the quality of life of patients and their families.

## Introduction

The ability to perceive hand actions of others plays a key role in our ability to learn fine motor skills from conspecifics and interact successfully with them in cooperative and competitive settings. C*erebral* cortical regions involved in motor control, including the premotor cortex and inferior parietal cortex where mirror neurons were found in the monkey (di Pellegrino *et al*., 1992; Gallese *et al*., 1996; Rizzolatti *et al*., 1996; Kohler *et al*., 2002; Keysers *et al*., 2003; Fogassi *et al*., 2005; Rozzi *et al*., 2008), and the primary somatosensory cortex (SI) (Gazzola and Keysers, 2009; Caspers *et al*., 2010; Keysers, Kaas and Gazzola, 2010), have been shown to be necessary for extracting subtle information from the observed kinematics of hand actions (Urgesi, Candidi and Avenanti, 2014; Keysers, Paracampo and Gazzola, 2018). A powerful task to reveal the impact of disturbing these cortical regions requires participants to judge the weight of an object lifted by another individual. This task depends on the ability to transform subtle kinematic cues into a weight estimate. Perturbing activity in the premotor cortex and SI disrupts the ability to perceive the weight (Pobric, De and Hamilton, 2006; Valchev *et al*., 2017), suggesting a causal role of premotor and somatosensory region in action perception.

The cerebellum is a key partner of these neocortical brain regions during motor control, where its role is well established (Kelly and Strick, 2003; Gao *et al*., 2018). It is perhaps not surprising that some have speculated that the cerebellum may also play a role in the perception and prediction of the kinematics of observed hand actions. Specifically, it has been proposed that the cerebellum could leverage its forward models (i.e. neural computations that transform motor signals into expected sensory consequences) to predict the actions of others (Miall, 2003; Wolpert, Doya and Kawato, 2003; Fuentes and Bastian, 2007; Gazzola and Keysers, 2009; Rizzolatti and Sinigaglia, 2010). Although this proposal is intuitively appealing, we still have little evidence for the cerebellum being a reliable and even necessary node of the action observation network (Sokolov, Miall and Ivry, 2017). This is because fMRI evidence for its recruitment during action observation is mixed, and very few neuro-modulation or lesion studies have explored the impact of cerebellar disruptions on hand action observation.

With a few exceptions, imaging studies on action perception have typically focused on the involvement of the neocortex, leaving the information about cerebellar activity limited to what the field of view of fMRI of these studies usually included, i.e. the dorsal cerebellum (e.g. (Aziz-Zadeh, 2006; Gazzola *et al*., 2007a; Gazzola *et al*., 2007b; Catmur *et al*., 2008; Gazzola and Keysers, 2009; Agnew, Wise and Leech, 2012; Brunner *et al*., 2014; Plata Bello *et al*., 2014; Di Cesare *et al*., 2015; Jelsone-Swain *et al*., 2015; Thomas *et al*., 2018)). Several other experimental studies fail to observe cerebellar activation to hand action observation (Iacoboni *et al*., 1999, 2001; Buccino *et al*., 2004; Orr *et al*., 2008; Rocca and Filippi, 2010; Jastorff, Abdollahi and Orban, 2012; Sasaki *et al*., 2012; Horan *et al*., 2014). This inconsistency is also reflected in meta-analyses of action observation studies, with some finding no (Caspers *et al*., 2010) or very limited cerebellar activations (Molenberghs, Cunnington and Mattingley, 2012), and others finding several clusters (Van Overwalle *et al*., 2014). In their extensive meta–analysis, Van Overwalle et al. found that only 28% of the reviewed studies investigating action observation report cerebellar activity. The degree to which these inconsistencies depend on data-acquisition and data-analysis pipelines not optimized for the cerebellum is difficult to estimate post-hoc, and experiments that optimize methods for the cerebellum, assess the reliability of activations in individual participants, and assess replicability across studies are required. Part I of the current manuscript will therefore present four fMRI experiments that map and replicate the recruitment of cerebellar voxels during hand action observation using MRI acquisition and analysis methods optimized for the cerebellum. These studies highlight that lobules VI and VIII are consistently recruited by action observation.

However, to establish whether the cerebellum causally contributes to hand action observation, its activity must be perturbed and the impact on action perception measured. Unfortunately, only two studies have taken that route so far. First, Sokolov (Sokolov *et al*., 2010) showed that four patients with tumours in the left lateral cerebellum (but not those with lesions in the vermis) were impaired in their ability to detect whether a point-light walking motion was embedded in random dot motion of that locomotor activity. However, the motor control of routine walking and of skilled hand actions is fundamentally different, as demonstrated by the fact that lesioning the pyramidal tract that transmits the cortical output to the spinal cord leaves routine treadmill walking unaltered (Eidelberg and Yu, 1981), but severely impairs skilled hand actions (Forssberg *et al*., 1999; Duque *et al*., 2003; Hermsdörfer *et al*., 2003). Second, Cattaneo (Cattaneo *et al*., 2012) tested the involvement of the cerebellum in the perception of action sequences. They showed eight participants affected by cerebellar ischemia sets of four still photographs taken during an action (e.g. opening a bottle and pouring a glass of water). One out of the four pictures was not fitting the temporal sequence of the action, and the task was to identify which one was the intruder. They found the performance of five of the cerebellar patients to be below the range of the sixteen healthy controls. While this study does not explore the processing of the subtle kinematic cues, it provides the first evidence that cerebellar impairments can affect the ability of participants to identify acts not belonging to a particular action sequence. However, while dozens of studies in hundreds of participants establish that premotor and parietal regions of the neocortex are necessary for the optimal perception of observed actions (Urgesi, Candidi and Avenanti, 2014; Keysers, Paracampo and Gazzola, 2018), the necessary role of the cerebellum in hand action observation hinges on a single study with 8 patients. In part II of the current study we therefore aim to provide new evidence for a contribution of the cerebellum to action perception, and the first evidence for its role in processing subtle kinematic cues during hand action perception. To this aim, we tested the ability of 21 patients with spinocerebellar ataxia of subtype 6 (SCA6) to detect the weight of a box by observing the kinematics of a hand lifting the box in a video setting. SCA6 is a rare late-onset neurodegenerative disorder characterized by ataxia and associated with a loss of Purkinje cells in the cerebellum. A Voxel-based morphology study points to loss of gray matter in the hemispheres of lobule VI (Rentiya *et al*., 2017) as being the primary cause of the upper limb ataxia – adjacent to regions in which we find cerebellar activations to action observation in part I. Task performance was compared with that of 31 age matched controls. Participants were tested in (i) a condition in which a sleeve on the actor’s arm occluded muscle shape information, forcing participants to focus on the arm’s kinematics to judge the weight of the box (Sleeve condition), and in (ii) a condition in which the sleeve was removed to reveal information on the appearance of muscle contractions, which complements the arm’s kinematic information (NoSleeve). Comparing the two groups in the Sleeve condition will reveal whether the cerebellum is necessary for kinematic processing. Comparing the gain in performance across the two conditions (i.e. the NoSleeve-Sleeve performance difference) across groups will reveal whether the cerebellum is necessary to extract additional information from biological shape.

The two main aims of our work are therefore to establish: (a) whether and where hand action observation reliably activates the cerebellum and (b) whether perturbations of cerebellar functioning impair the ability to process the kinematic and/or shape of observed actions.

## Materials and Methods

### General overview of the experiments and participants)Table 1)

Experiment #1 was aimed at localizing cerebellar activity to action observation using different analysis pipelines, and at comparing the results between pipelines and to those found in the literature. Experiment #2 and #3 tested the replicability of the results of Experiment #1 on two independent samples of participants, and on a different MRI scanner. Experiment #4 tested the impact of the weight discrimination task on the previously identified action observation network, and Experiment#5 was aimed at directly testing the involvement of the cerebellum in action perception by comparing the accuracy in weight estimation between SCA6 patients and matched controls.

**Table 1.**
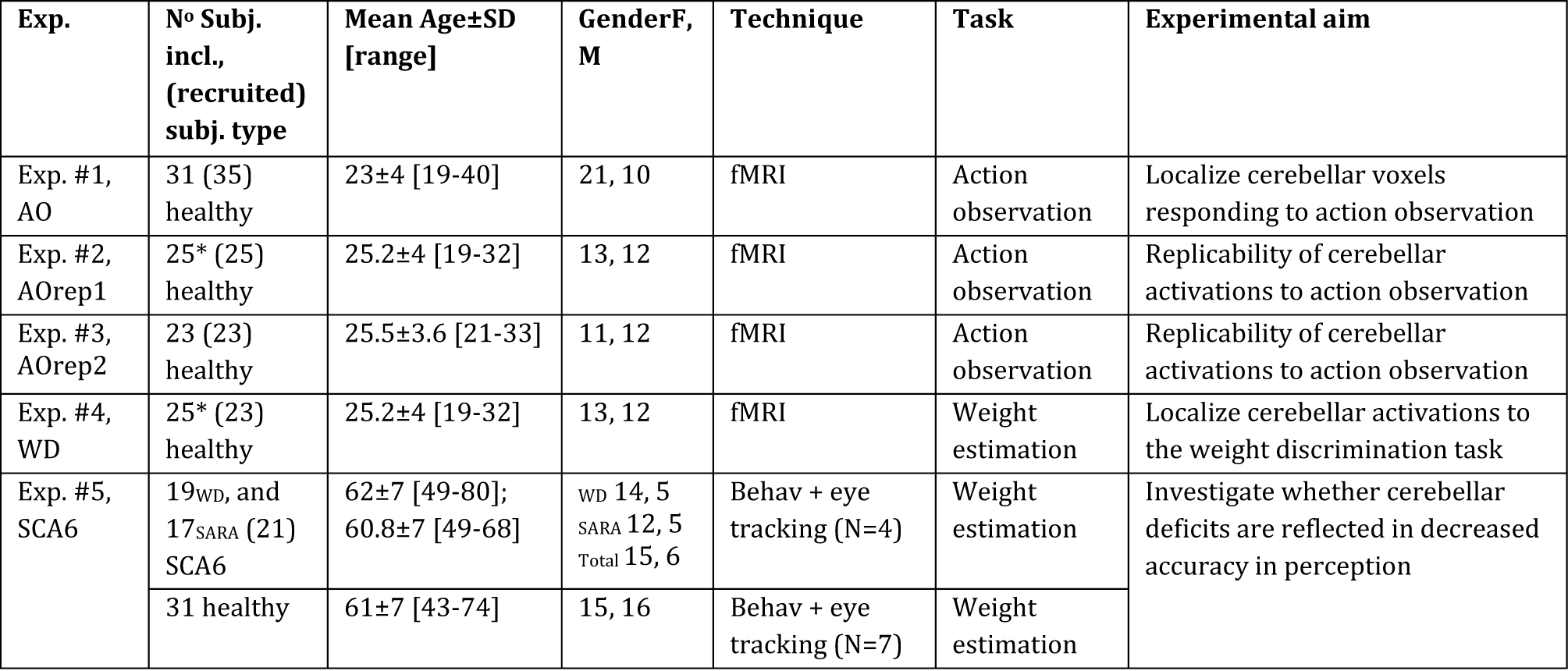
Experiments overview. From left to right: the acronym of each experiment (AO=action observation; AOrep=action observation replication; SCA6=spinocerebellar ataxia type 6; WD=weight discrimination; SARA= Scale of the Assessment and Rating of Ataxia); the number of participants included in the analyses and their characterization (number in brackets indicate the number of initially recruited participants); the average age of the group, its standard deviation and range in brackets; the number of females and males within each group of included participants; the technique involved in the experiment; the task used for each experiment; the aim of each experiment. All groups of participants are independent except the ones marked with *, in which the same 25 participants underwent both the passive observation and the weight estimation task in separate sessions. In Exp. #1, four participants were excluded from the statistical analysis: two due to excessive head motion (displacement of more than the 3.5 mm voxel dimension), one reported sleepiness, and one because of image distortion. In Exp. #5, two participants were excluded from the weight lifting task because pre-symptomatic, and two more were excluded from the correlation with SARA because did not have SARA scores.

| Exp. | N° Subj. incl., (recruited) subj. type | Mean Age±SD [range] | GenderF, M | Technique | Task | Experimental aim |
| --- | --- | --- | --- | --- | --- | --- |
| Exp. #1, AO | 31 (35) healthy | 23±4 [19-40] | 21, 10 | fMRI | Action observation | Localize cerebellar voxels responding to action observation |
| Exp. #2, AOrep1 | 25* (25) healthy | 25.2±4 [19-32] | 13, 12 | fMRI | Action observation | Replicability of cerebellar activations to action observation |
| Exp. #3, AOrep2 | 23 (23) healthy | 25.5±3.6 [21-33] | 11, 12 | fMRI | Action observation | Replicability of cerebellar activations to action observation |
| Exp. #4, WD | 25* (23) healthy | 25.2±4 [19-32] | 13, 12 | fMRI | Weight estimation | Localize cerebellar activations to the weight discrimination task |
| Exp. #5, SCA6 | 19<sub>WD</sub>, and 17<sub>SARA</sub> (21) SCA6 | 62±7 [49-80]; 60.8±7 [49-68] | WD 14, 5<br>SARA 12, 5<br>Total 15, 6 | Behav + eye tracking (N=4) | Weight estimation | Investigate whether cerebellar deficits are reflected in decreased accuracy in perception |
| | 31 healthy | 61±7 [43-74] | 15, 16 | Behav + eye tracking (N=7) | Weight estimation | |

All tested healthy participants had a normal or corrected to normal vision, and none had a history of neurological conditions or treatments. The participants tested in the MRI also met MRI safety requirements.

The SCA6 patient group was recruited in collaboration with the department of Neurology at the Erasmus MC Rotterdam (see Supplementary method 1.0). The severity of disease progression was clinically assessed by a licensed neurologist using the Scale of the Assessment and Rating of Ataxia (SARA) (Schmitz-Hubsch *et al*., 2006; Saute *et al*., 2012). SARA includes 8 items (gait, stance, sitting, speech disturbance, finger chase, nose-finger test, fast alternating hand movements and heel-shin slide) reflecting neurological manifestations of cerebellar ataxia (Weyer *et al*., 2007). SARA scores range from 0 to 40, with higher scores corresponding to higher progression. The average SARA score for our patients group (N_SARA_=17) was 11.38 ± 5.75 (SD; range: 2 to 21.5). The thirty-one healthy participants that were recruited as control group, matched the SCA6 group for age (t _(50)_=0.96, p=0.34), handedness (SCA6: 19 right handed and 2 left handed, Controls: 27 right and 4 left handed, Yates corrected X^2^=0, p=0.94) and gender (SCA6 15f:6m, ctrl 15f:16m, Yates corrected X^2^=1.86, p=0.17). However, our patient group contained numerically fewer males, an issue that is addressed in control analyses. Controls did not receive a clinical assessment.

All participants signed an informed consent in accordance with the declaration of Helsinki. The fMRI study protocols were approved by the medical ethical committee of the University of Groningen (METc2012/380), the ethics review board of the University of Amsterdam (2015-BC-4697), the Academic Medical Center of Amsterdam (W15_243#15.0288), and the clinical study protocol was approved by the Medical Ethical Committee of the Erasmus MC Rotterdam (MEC-2013-095).

### Stimuli, tasks and paradigms

#### Action observation task (Fig. 1A)

During the observation task participants watched 39 unique movies of a human right hand interacting with objects displayed on a table (ActionOBS). The 39 control movies displayed a hand movement without a meaningful object interaction (CtrlOBS). Exp. #1 and 2 also contained a third static condition, in which the hand rested close to the object (stimuli also shown in Arnstein *et al*., 2011; Valchev *et al*., 2016). This static condition was not included in Exp. 3, and therefore not included in the group analyses. Conditions were randomized across participants and presented using the Presentation^®^ software (Version 18.0, Neurobehavioral Systems, Inc., Berkeley, CA, www.neurobs.com) in a single fMRI run. Participants were instructed to pay close attention to the movies shown.

**Fig. 1.**
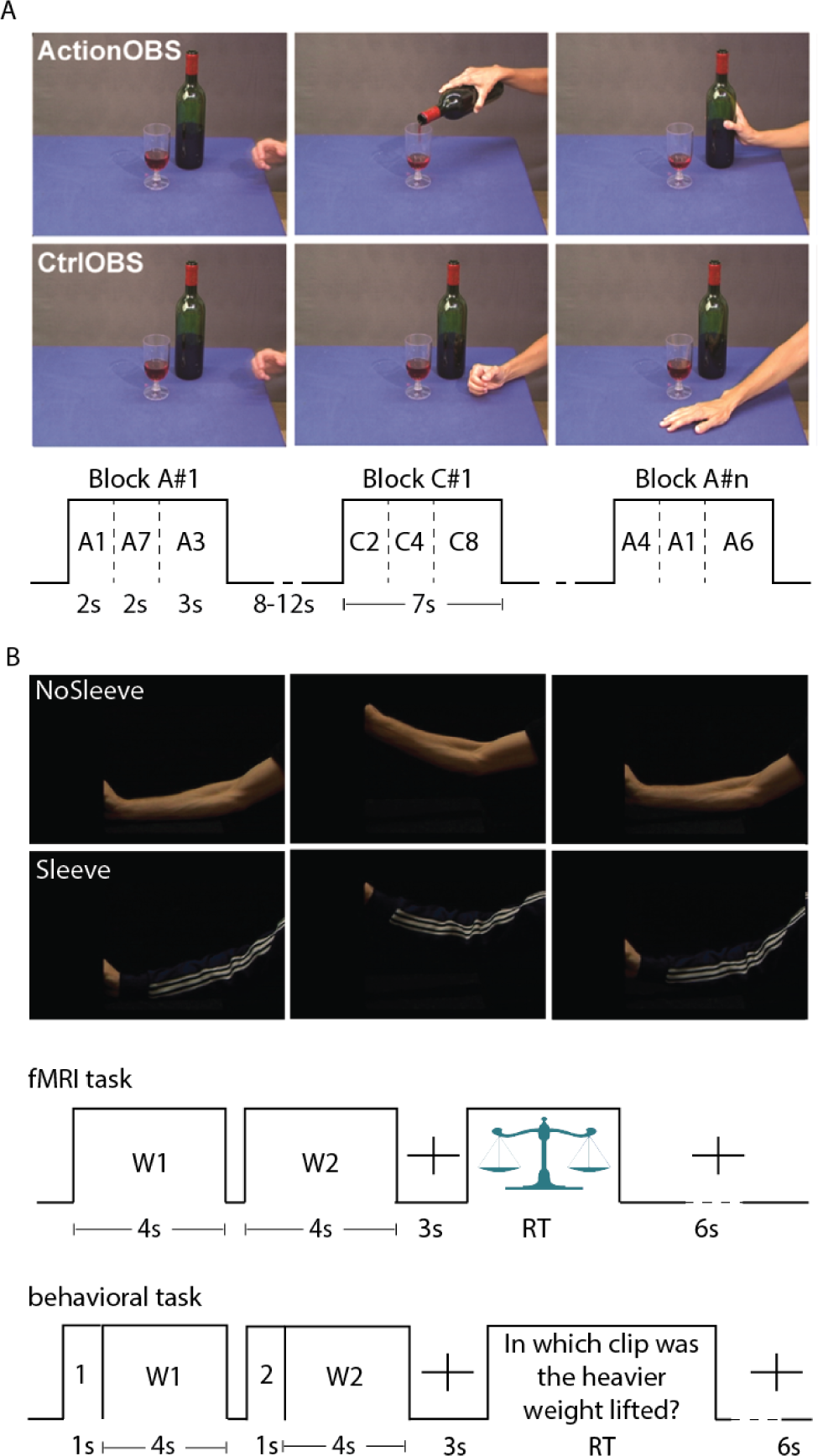
Experimental tasks. **(A) Action observation task.** Example of one out of the 39 possible actions and its control, followed by the task structure. Ctrl= control. OBS=observation. A=action. C=control. The ActionOBS and CtrlOBS movies were grouped in blocks of 7 seconds. Each block contained three actions from the same condition, with a total of 13 blocks for each condition. Blocks were separated by a fixation cross for a random period of 8 to 12 seconds, displayed on a background that was visually similar to the table. **(B) Weight discrimination task.** Frame extracted from the NoSleeve (top) and Sleeve (bottom) weight lifting condition, followed by the trial structure for the fMRI (top) and behavioral experiments (bottom). In the fMRI task the window of time participants were requested to answer was indicated by a weighing scale. In the behavioural task clips were preceded by the number 1 or 2 denoting whether it was the first or second clip of the pair. The sentence following the video was translated from Dutch for illustration purposes. RT=participant’s reaction time.

#### Weight discrimination task (WD, Fig. 1B)

Participants performed a two-alternative forced-choice task, in which at every trial, participants had to choose in which of the two presented videos the heaviest object was lifted. The 4 seconds video-clips showed a human arm lifting an object. In order to avoid participants to deduce the weight from object movement only (e.g. differences in object shaking during the lifting phase), a black panel occluded both the object and the hand from vision. To disentangle whether the contribution of cerebellum mainly comes from computation of action kinematics or from arm shape information, two versions of the task were created: (i) in half of the trials, the arm lifting the object was sleeved thus making the kinematic of the arm the only information available to perform the task (Sleeve); (ii) in the other half, the arm was uncovered thus allowing both kinematic and shape information to be used (NoSleeve; Fig. 1B). During the video recording, the actor was instructed to lift one of three weights (2850g, 900g and 180g) within 4 seconds. A metronome was used to time the lift, and a reference line was marked on the wall in front of the actor to help maintaining the same lifting height throughout all videos. The actor was aware of the object weight to avoid hesitation in the lifting. Videos were recorded using a digital video camera (Sony DSRPDX10P) and edited using Adobe Premiere Pro (Version CS5, Adobe System Incorporated, San Jose, USA). Clips showing the same lifted weight were never paired. In half of the trials the heaviest object was lifted first, in the other half as second. The order was randomized in Psychopy2 (Peirce, 2009). After the second clip, the task instruction was presented until the subject indicated his/her response. Before the beginning of the task participants performed four training trials.

Some minor task differences were present between Exp#4 and #5 (Fig. 1B).

#### Exp. #5 (behaviour)

Participants gave the response by pressing the arrow keys on a standard QWERTY keyboard using their right hand. Ninety-six trials were presented in total, and participants had the option to take a short voluntary break after the first half of the trials.

#### Exp. #4 (fMRI)

Participants indicated their responses by means of a MRI compatible button box. Participant used their left hand to select the first clip and their right hand to select the second. Stimuli were presented using Presentation^®^ software. For the fMRI experiment, a numerosity task was additionally introduced and intermixed with the weight discrimination task. Participants had to estimate and compare the number of moving dots shown in video 1 and 2 instead of weight. The movement of the dots followed the kinematic of the arm presented in the Sleeve and NoSleeve conditions, but the arm was not visible. Since an error occurred in the randomization of this condition, and this task was not performed by the SCA6 group, the numerosity condition was not included in the group analyses. Seventy-two trials were presented in total (24 for each of the three conditions).

### fMRI data acquisition

All MRI data sets included an anatomical scan. Exp. #1 then included one functional scan of the action observation task. Exp. #2 and 3 aimed at comparing the effect of different numbers of simultaneous slice acquisition on task based fMRI, and included four and five functional scans of action observation respectively. The results of this comparison are the subject of a separate manuscript. Because participants of Exp. #1 only saw the movies once, we only included the first view of the action observation task, independently of the number of simultaneously acquired slices. Exp. #4 included two functional runs of the weight lifting task. These two runs were randomly presented between the four observation runs of Exp. #2. The scanning parameters were chosen to achieve a coverage of the entire cerebrum and cerebellum (Supplementary Table 1).

### Localization of cerebellar activations, impact of different analysis pipelines and replicability

The impact of different pipelines on cerebellar task-based responses was analysed on data from Exp. #1. The four considered pipelines mainly differed in the order in which the pre-processing and first level subject statistics were computed, and in the normalization template. Because the comparison revealed a no clear advantage of using pipelines optimized for the cerebellum compared to the traditional one, the method and results of this comparison are presented in the supplementary material.

All the analyses included in the main text therefore follow the traditional approach which includes: slice-time correction, realignment of functional images to the computed mean, co-registration of the anatomical image to the mean, whole brain normalization to the MNI template (final voxel size: 2 × 2 × 2 mm) based on the parameter generated during the segmentation of the co-register anatomy, a smoothing with a 6 mm FMHW Gaussian kernel followed by a general linear model (GLM). Analyses testing the possibility of activation leakage between the anterior cerebellum and the temporal cortex due to smoothing are reported in the supplementary material.

For Exp.#1 to 3, the GLM included two standard box car predictors that modelled the ActionOBS and CtrlOBS video presentation. Exp. #1-2 also included a predictor modelling the static conditions. All predictors were convolved with the canonical hemodynamic response function (HRF). The last six regressors of no interest included the displacements and rotations along the three axes, determined during image realignment. The ActionOBS-CtrlOBS contrast was computed at the subject-level to generate action specific activations for observation. Analyses of variance on the ActionOBS-CtrlOBS contrast values from Exp. #1-3 were also implemented to directly compare the results of the three experiments to each other (within-subjects ANOVA) as well as to baseline (one-way ANOVA).

All analyses were run in SPM8 and 12 (Wellcome Trust Centre for Neuroimaging, UCL, UK) using Matlab 7.14 (The MathWorks Inc., Natick, USA) with a bounding box size adjusted to include the entire cerebellum [−90 −126 −72; 91 91 109], complemented by custom Matlab scripts. Unless specified otherwise, all analyses were estimated within the cerebellar mask using the cerebellar anatomical map from the Anatomy toolbox (http://www.fz-juelich.de/ime/spm_anatomy_toolbox) (Geyer *et al*., 1996, 2000; Amunts *et al*., 1999; Geyer, Schleicher and Zilles, 1999; Grefkes *et al*., 2001; Geyer, 2004; Eickhoff *et al*., 2005, 2006, 2007; Caspers *et al*., 2006; Choi *et al*., 2006). The Anatomy toolbox was also used to define regions of interests, and guide anatomical descriptions of clusters of activity.

Unless specified otherwise, all statistical maps were thresholded at p_*FWE*_<0.05 with a minimal cluster size of 10 voxels. We chose peak-level FWE-correction, because we wished to (i) interpret activation of individual voxels, and, motivated by the inconsistencies of cerebellar activations in the literature, (ii) to limit the risks of Type I errors.

In order to investigate the consistency in location of voxels responding to action observation between participants and studies, we computed consistency maps (Gazzola and Keysers, 2009, and Supplementary method 1.2). However, as the consistency maps cannot confirm that voxels responding to action observations are present in all participants, we counted the number of activated voxels within each participant. This counting was done separately for the four cerebellar anatomical regions of interest (left and right lobule VI, and VIIb/VIIIa), and for the cerebellum as a whole. To compare the reliability of cerebellar activations with that of the cortex, the counting was done for three additional cortical regions, typically associated with the action observation network (Gazzola and Keysers, 2009; Caspers *et al*., 2010; Molenberghs, Cunnington and Mattingley, 2012): the premotor area BA44, the inferior parietal complex PF and the primary somatosensory cortex SI.

### Localization of the weight discrimination task

The GLM of Exp. #4 included eight boxcar predictors: three modelled the video presentation (i.e. from the beginning of Video 1 to the end of Video 2) associated to the Sleeve, NoSleeve and Numerosity conditions; two captured the participants responses at the time the weighting scale was presented separately for the left and right hand; one captured text information given to our participants at the beginning and the end of the each session; one included button presses that happened outside the response window; and one included the four videos used for training (only for the first session). The six head motion parameters were again added as covariate of no interest. Analyses of variance were used to compare the Sleeve and NoSleeve conditions to each other (within-subjects ANOVA), and to baseline (one-way ANOVA). As for Exp.#1-3, unless specified otherwise, the ANOVAs were computed within the cerebellar mask, at p*FWE<0*.*05*.

To test whether the videos used for the weight estimation task elicited activity in the areas to be found active for general action observation, an additional GLM was computed within a binary mask obtained by the global null conjunction of Exp. #1, 2 and #3 [Exp#1_ActionOBS-CtrlOBS_ OR Exp#2_ActionOBS-CtrlOBS_ OR Exp#3_ActionOBS-CtrlOBS_] (t_FWE_=2.06) from the one-way ANOVAs that included the ActionOBS-CtrlOBS from all three experiments. Results are shown at p_*FWE*_ <0.05.

### Analyses of behavioral data

Task performance scores were calculated as proportion of correct responses. We checked their normality using the Lilliefors test. Performance for the Sleeve and for the NoSleeve-Sleeve difference were normally distributed (both p>0.12). The performance in the NoSleeve condition and the average score of Sleeve and NoSleeve violated normality (both p<0.002). Accordingly, we used nonparametric tests as our main approach, and parametric analyses (ANOVAs and Bayesian analyses) were only used to supplement analyses for the Sleeve and NoSleeve-Sleeve difference.

To make sure the deficits in action perception did not occur because of visual tracking problems, eye tracking data were collected from four patients and seven healthy subjects; these control data as well as the methods for eye tracking are presented in the supplementary material.

## Results

### Localization of Action Observation Activations in the Cerebellum and their reliability

Viewing goal directed hand actions compared to control stimuli (ActionOBS-CtrlOBS) in Exp. #1 bilaterally recruited cerebellar Lobules VI, VIIb and VIIIa (Table 2, Fig. 2A and S1).

**Table 2.**
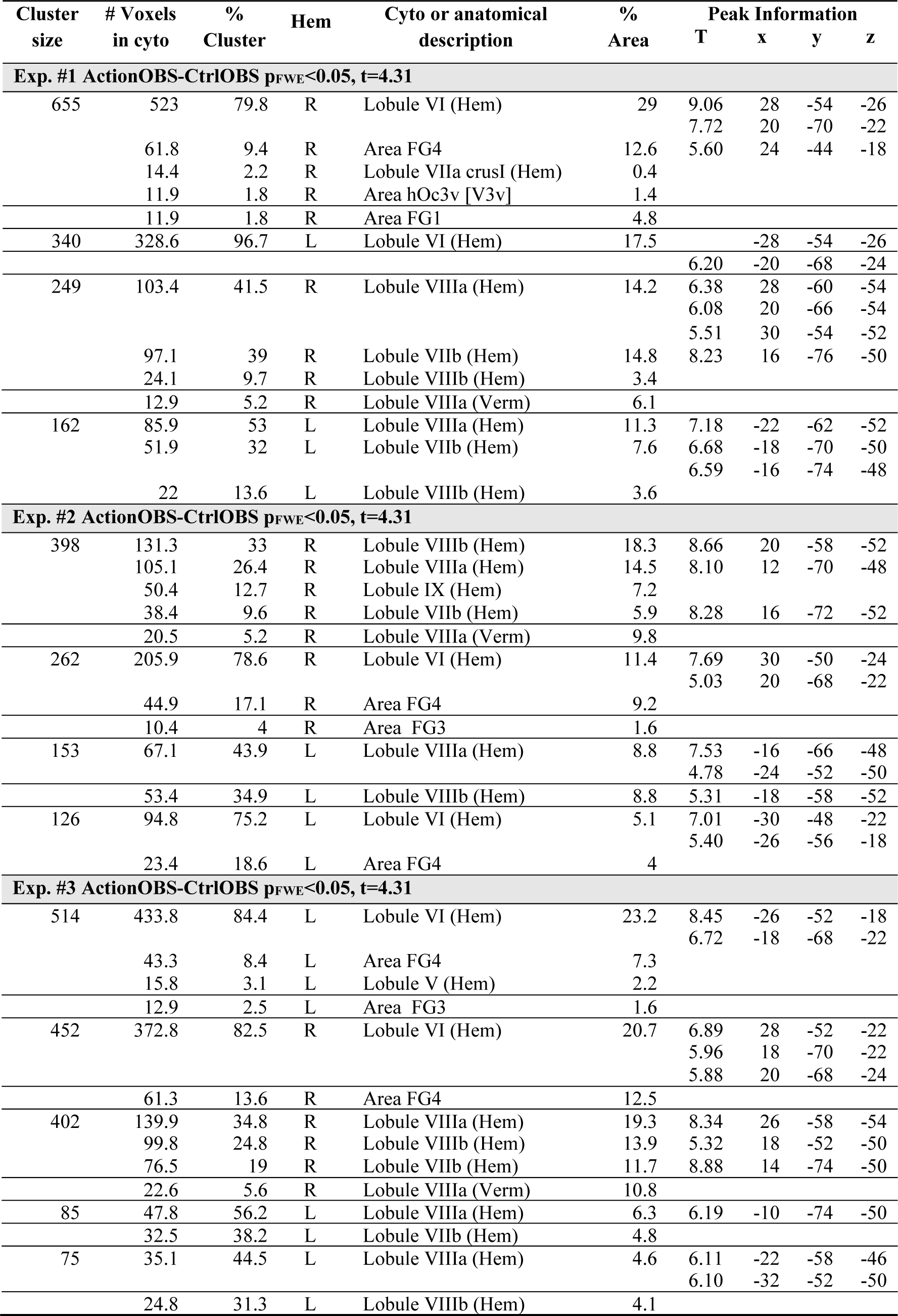
Cerebellar activations to ActionOBS-CtrlOBS for Exp. #1 to #3. Regions with ActionOBS-CtrlOBS≥4.31 labelled using SPM Anatomy Toolbox. Results are shown at p_FWE_ < 0.05 with cluster size >10 voxels. From left to right: the cluster size in number of voxels, the number of voxels falling in a cyto-architectonic area, the percentage of the cluster that falls in the cyto-architectonic area, the hemisphere (L=left; R=right), the name of the cyto-architectonic area when available or the anatomical description, the percentage of the area that is activated by the cluster, the t values of the peaks associated with the cluster followed by their MNI coordinates in mm.

**Fig. 2.**
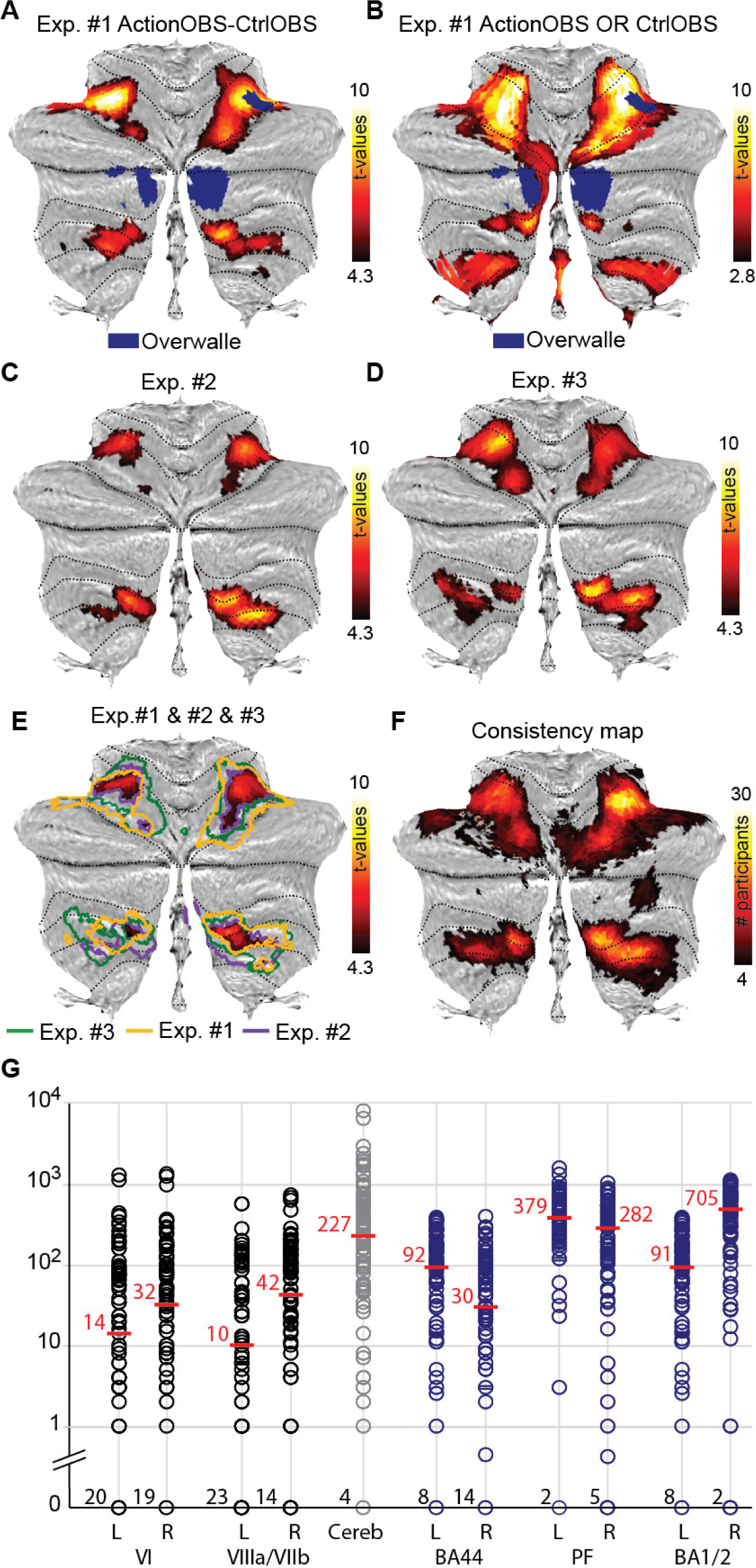
Reliability of cerebellar action observation activations. **(A-B).** In blue the maps presented by Van Overwalle and colleagues in 2014, and the results of the ActionOBS-CtrlOBS contrast of Exp. #1 in the hot color scale in (A) and of the global null conjunction ActionOBS OR CtrlOBS for Exp. #1 in (B), both at p_FWE_<0.05. **(C-D)** ActioOBS-CtrlOBS related activity for Exp. #2 and #3 respectively. p_FWE_<0.05, t=4.3. **(E)** Activations common to Exp. #1-3. Yellow, blue and green contours indicate the borders of the clusters shown in A, C and D to facilitate the qualitative comparison. **(F)** Consistency map computed on the smoothed data for the ActionOBS-CtrlOBS (p_unc_<0.001, t=3.1) contrast across the three experiments. The hot scale indicates the number of participant for which a particular voxel was significantly activated by the ActionOBS-CtrlOBS contrast. **(G)** Circles indicate the number of voxel a given subject had in each of the four cerebellar clusters (first four columns, black circles), in total in the cerebellum (fifth column, gray circles), and in three cortical regions also commonly activated by the ActionOBS-CtrlOBS contrast. The median is indicated by the red lines and numbers. Data are presented on a logarithmic scale and the number of participants having no voxels in a particular cluster is indicated on the x-axis.

Overlapping our activations with action observation maps from the meta-analysis of Van Overwalle and colleagues (Van Overwalle *et al*., 2014) (blue clusters of Fig. 2A, and Supplementary results 2.1) reveals only a small portion of the right lobule VI is common among the two maps. To test whether the limited overlap is due to subtracting our control condition, we overlapped the meta-analysis map with a global null conjunction of our conditions (i.e. ActionOBS OR CtrlOBS, p_FWE_<0.05, t=2.8). The overlap remained limited to right Lobule VI (Fig. 2B).

Considering this inconsistency, we (1) replicated the experiment on a different scanner in two new groups of participants and (2) explored how many of our participants had activations in the cerebellum.

Replicating the analysis in new participants confirmed the cerebellar recruitment, despite differences in scanning location and parameters (Fig. 2C-E, Table 2-3).

**Table 3.**
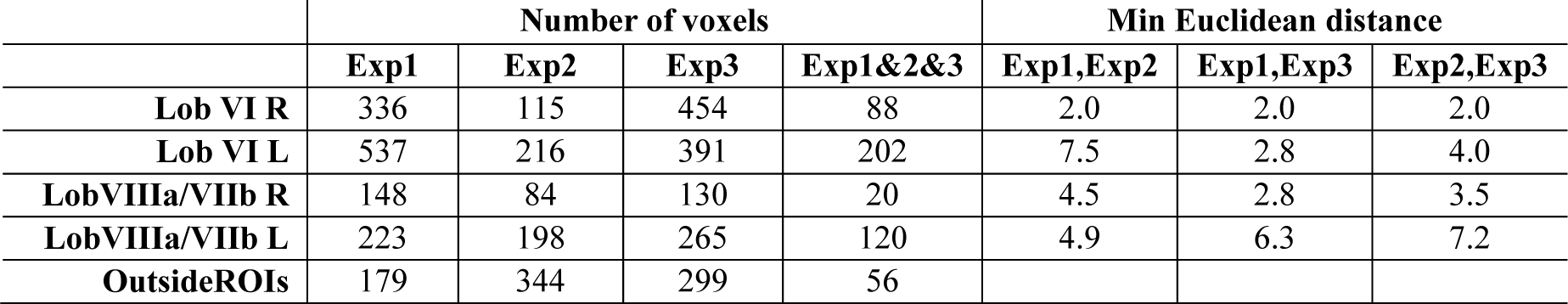
Comparison between Exp #1, #2 amd #3 in number of voxel and peak distance per cluster of activity. For each of the four cerebellar clusters, and for each experiment separately, the number of voxels surviving p_FWE_<0.05 for the contrast ActionOBS-CtrlOBS is reported. The fourth column reports the number of voxels counted within the conjunction of the three experiments. The last row indicates the number of cerebellar voxels not falling within the region of interest. Columns 5-7 indicate the minimum Euclidean distance between the activation-peaks identified belonging to the four clusters by the Anatomy toolbox for SPM.

Looking at individual participants revealed that all but four (all from Exp. #1) of the 79 participants had significant activations to the ActionOBS-CtrlOBS contrast when tested at p<0.001 (t=3.1) within the cerebellum (blue circles in Fig. 2G). The majority (68/79, 86.1%) additionally had >10 voxels activated (Fig. 2G and first 5 columns of Supplementary Table 4) and most had at least 10 voxels in each of the cerebellar lobules identified in the group (ROIs encompassing lobule VI or lobule VIIb+VIIIa). A binomial distribution indicates that finding 10 or more voxels significant by chance at p=0.001 in a ROI of 2085 voxels (the largest ROI we have) is highly unlikely (p<2×10^−5^).

To compare the reliability of cerebellar activations with those of the cerebrum, we took three regions consistently associated with the action observation system, BA44, the PF complex and SI (Keysers and Gazzola, 2009; Caspers *et al*., 2010; Molenberghs, Cunnington and Mattingley, 2012), and counted activated voxels in these regions subject by subject (Supplementary Table 4, last 6 columns). Chi^2^ tests comparing the proportion of participants with zero voxels activated in the 4 cerebellar and 6 cerebral regions using Fisher’s exact test in R indicated that for Exp#1 and #2 the proportion with zero voxels activated was larger in the cerebellum (Exp1, p=0.001; Exp2, p=0.004; Exp3, p=0.86). When combining all three experiments, the difference in proportion became highly significant (p<0.001), with the cerebral ROIs hosting significant voxels in a larger proportion of participants than the cerebellar ROIs.

Consistency maps indicated that the right Lobule VI hosted the most consistently activated voxel with 30 participants having significant activations in that specific voxel (Fig. 2F).

In summary we find that our task reliably activates the cerebellum at the individual and group level, and across scanning location and pipelines. Despite the high reliability of our task the results however only overlap with those in the literature in a small part of right Lobule VI, and remain less reliable than cerebral activations.

### Cerebellar activation to the weight discrimination task

Observing an arm lifting an object to judge its weight activates several regions of the cerebellum (Fig. 3A,B and Table 4, p_FWE_<0.05, t=2.8). The responses to lifting movements overlap with the ALE-meta-analysis maps (Van Overwalle *et al*., 2014) beyond Lobule VI, in bilateral Lobule VIIa crusI. Computing the GLM of the weight discrimination experiment within the global null mask of the previous three experiments shows that all clusters observed in Exp. #1-3 were activated by the observation of lifting movement (Fig. 3C,D, Table 4).

**Fig. 3.**
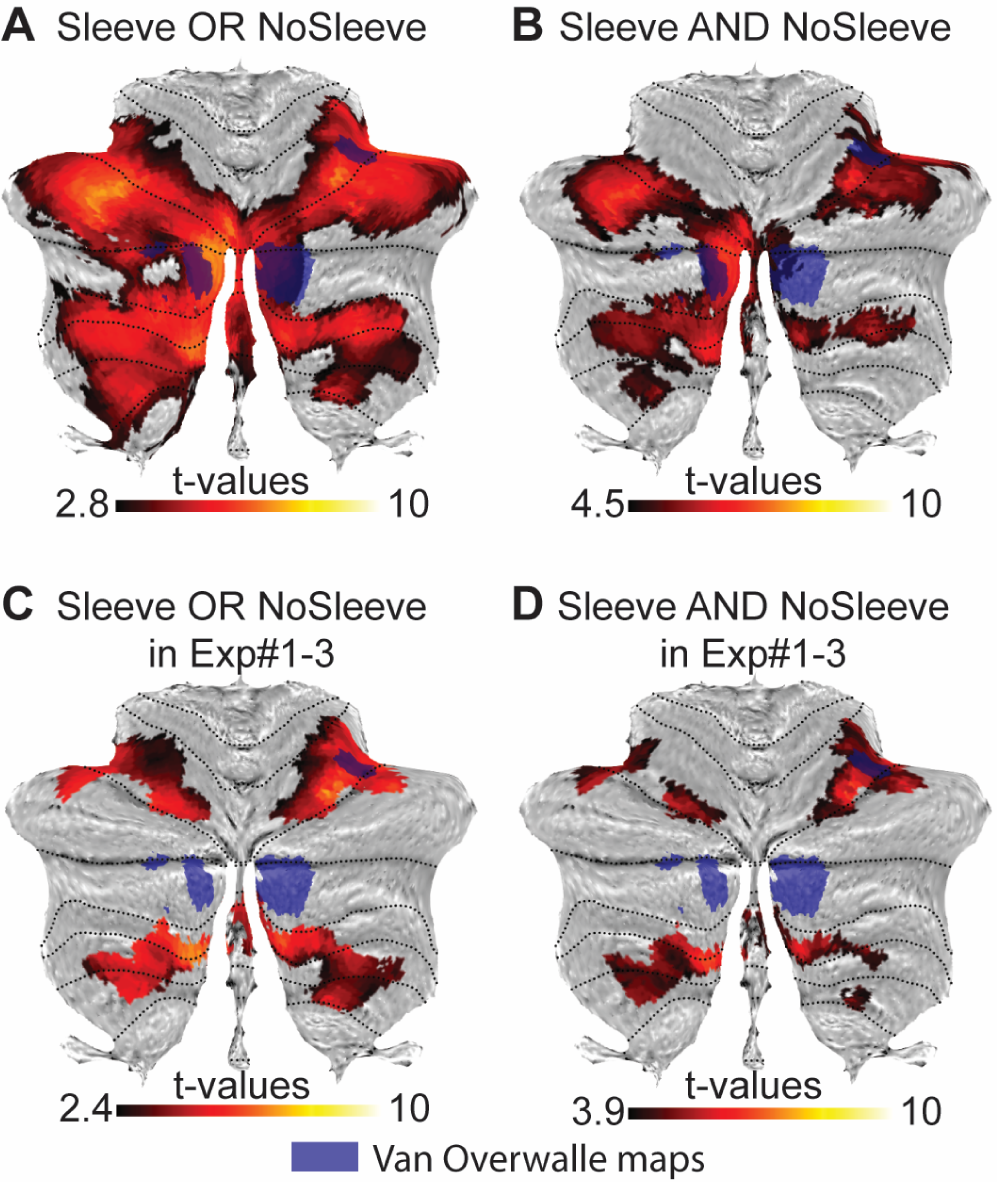
fMRI results of the weight discrimination task. **(A)** Voxels significantly activated by either the Sleeve (only kinematic information available) or the NoSleeve (both kinematic and shape information) condition (global null conjunction in SPM at p_FWE_<0.05, t=2.8, min 10 vx). In blue the clusters identified by Van Overwalle et al. 2014, as responding to action perception. **(B)** Voxels activated by both (conjunction-conjunction in SPM) by the NoSleeve and Sleeve conditions (p_FWE_<0.05; t=4.5, min 10 voxels). **(C)** Same as in (A) but within the clusters of activation found in Exp. #1 to #3 (Exp.#1>0 OR Exp.#2>0 OR Exp.#3>0). Results are shown at p_FWE_<0.05, t=2.8, min 10 voxel. **(D)** Same as in (C) but within the clusters of activation found in Exp. #1 to #3 (p_FWE_<0.05; t=3.9, min 10 voxels). All activations are shown on the flat map of the cerebellum offered by the SUIT toolbox.

**Table 4.**
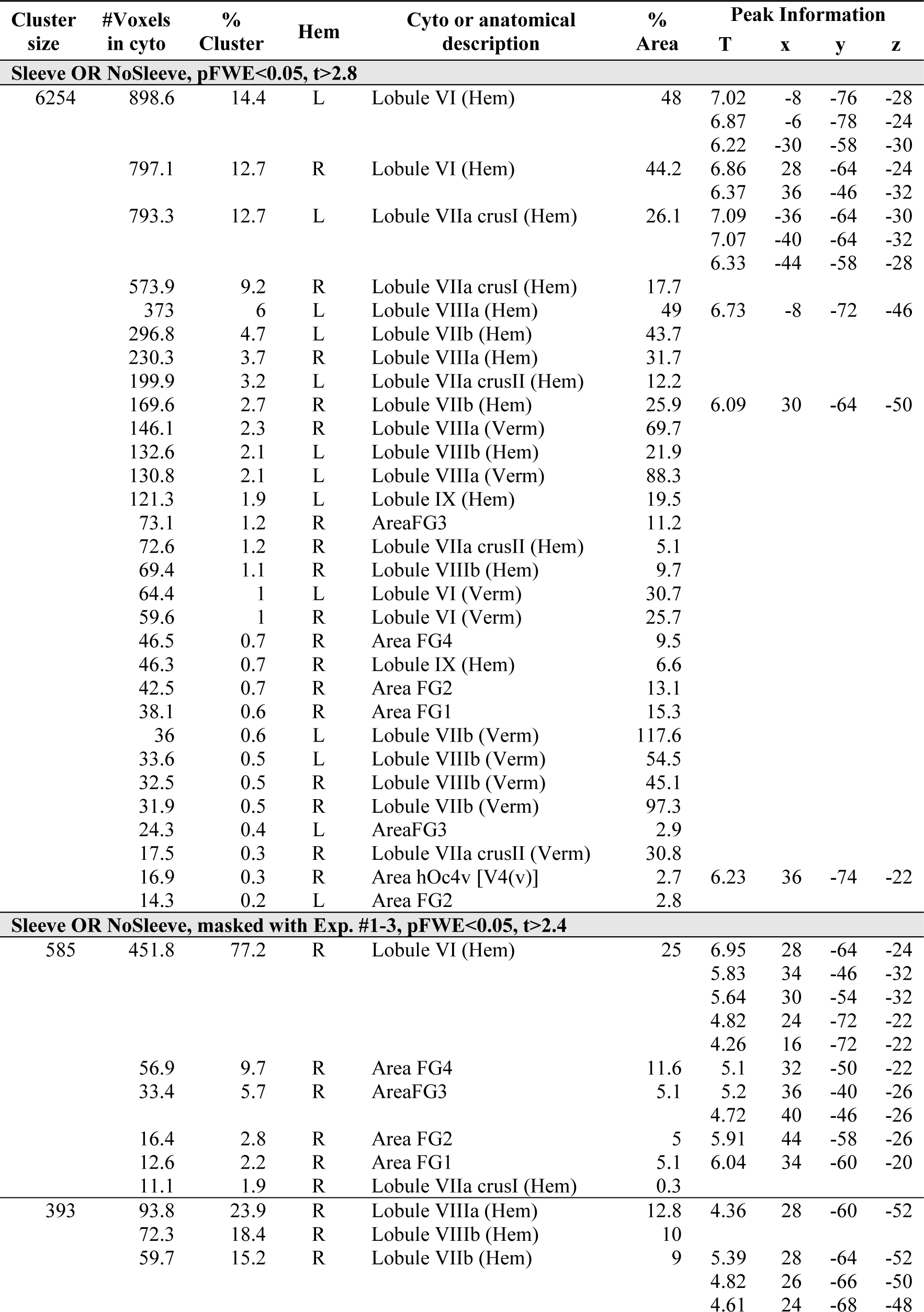
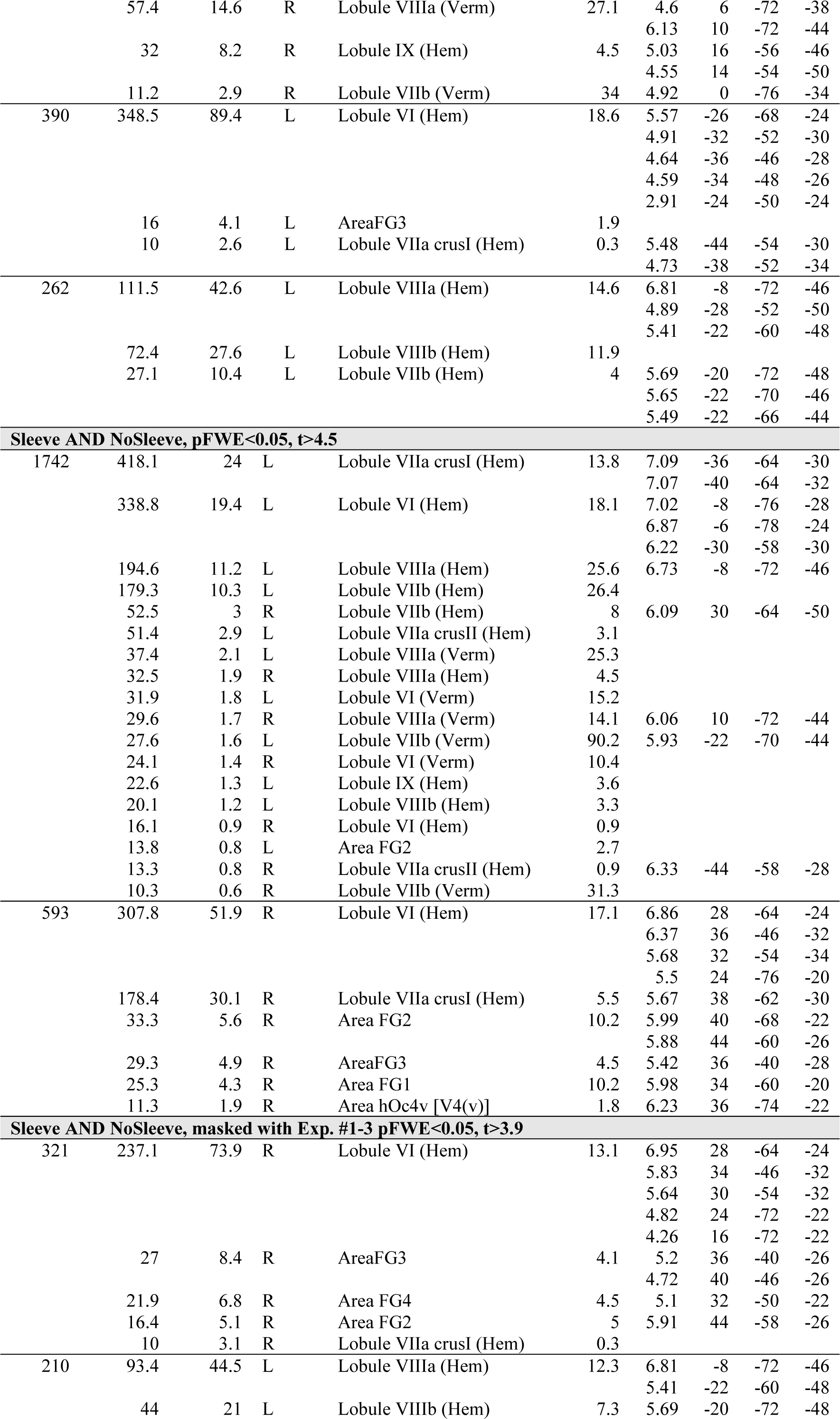
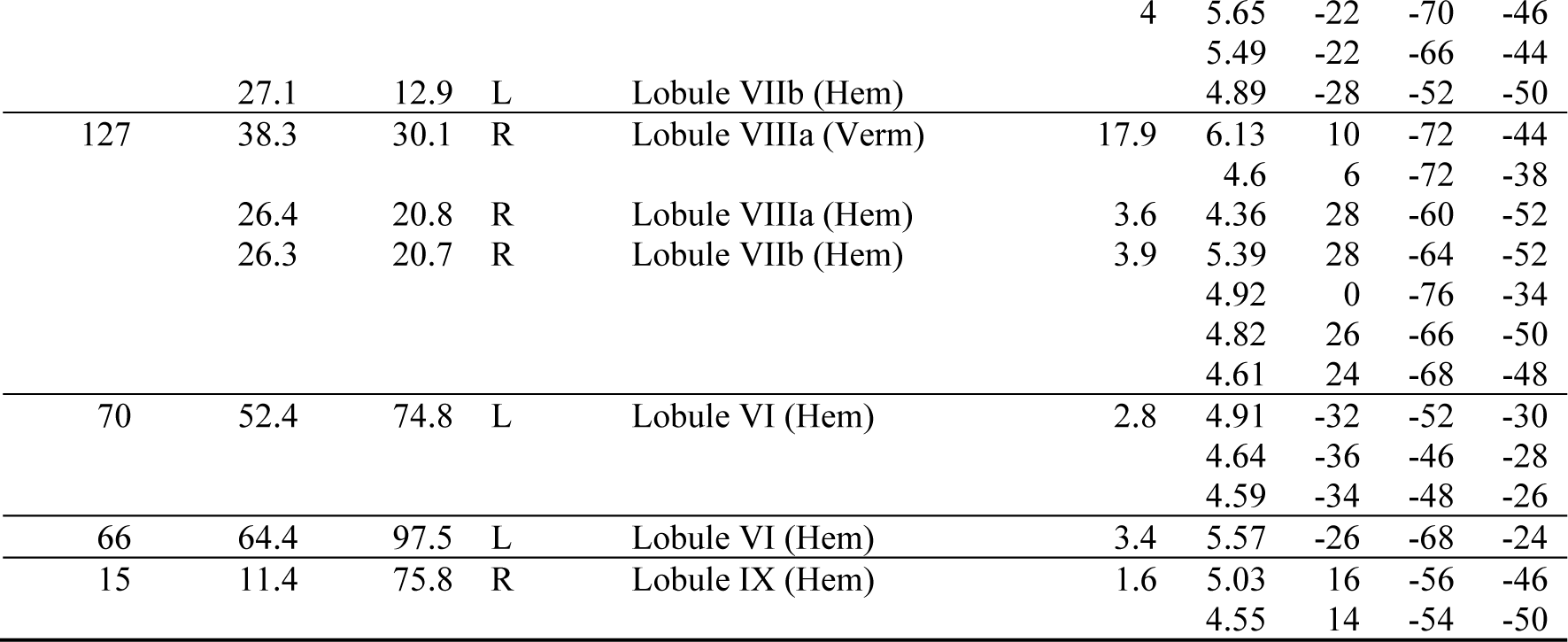
Cerebellar activations to the weight discrimination task. Results are shown at p_FWE_<0.05 with cluster size >10 voxels. Conventions as in Table 3.

**Sleeve OR NoSleeve, masked with Exp. #1-3, pFWE<0.05, t>2.4**

**Sleeve AND NoSleeve, pFWE<0.05, t>4.5**

What aspect of action observation is processed in the cerebellum? By disentangling the activity common to the Sleeve and NoSleeve conditions mentioned above (Conjunction Sleeve & NoSleeve) from that specific to the NoSleeve condition (NoSleeve-Sleeve), we can attempt to identify regions involved in kinematic and shape processing, respectively. The eye-tracking maps from the control participants show that the two conditions are indeed explored differently (Supplementary Fig. 2A). When the arm was covered participants focussed similarly on the proximal and distal part of the arm (t_(12)_=1.523, p=0.154) but if no sleeve was present participants focused significantly more on the proximal part of the arm (t_(12)_=-9.482, p<0.001) that reveals shape information in the upper arm musculature. Results from the fMRI data indicate that in contrast to the conjunction that revealed consistent cerebellar involvement for kinematic processing, at FWE correction at peak level nothing survive for both the Sleeve-NoSleeve and the NoSleeve-Sleeve contrast within the cerebellum (t=4.42, p>0.05), while 22 voxels in the fusiform area FG4 become apparent for the contrast NoSleeve-Sleeve when the analyses is run for the whole brain (t=5.4, p<0.05). Accordingly, the cerebellum is significantly recruited by the kinematic cues common to both conditions (Fig. 3), but not by the differential shape cue that the NoSleeve-Sleeve contrast situates in the ventral visual stream instead.

### Cerebellar contribution to action perception

The Mann-Whitney U test on task performance revealed a significant difference between SCA6 and controls for the Sleeve condition (N_SCA6_=21; N_ctrl_=31; U=199.5; p<0.009), in which participants depend on the kinematic information (Fig. 4A). The same test revealed that the gain of performance in the NoSleeve compared to the Sleeve condition (i.e. NoSleeve performance – Sleeve performance) did not differ significantly across groups (N_SCA6_=21; N_ctrl_=31; U=274.5; p>0.34). Not surprisingly, the two groups therefore also differed when the total performance was considered, including both the Sleeve and NoSleeve trials (N_SCA6_=21; N_ctrl_=31; U=183; p<0.004). Using d’ instead of percent correct led to similar conclusions. To explore whether our pattern of findings, which included a significant group difference for the Sleeve condition and a lack of significant group difference in the gain of performance, was evidence that the cerebellum contributes to kinematic but not shape processing in our experiment, we performed a Bayesian t-test in JASP. The Bayes factors in favour of the alternative hypothesis Ctrl>SCA6 were BF=14.7 (Sleeve) and BF=0.19 (NoSleeve-Sleeve performance). Accordingly, we have strong evidence for a group difference in kinematic processing (Sleeve), and moderate evidence for a lack of difference for shape processing (NoSleeve-Sleeve).

**Fig. 4.**
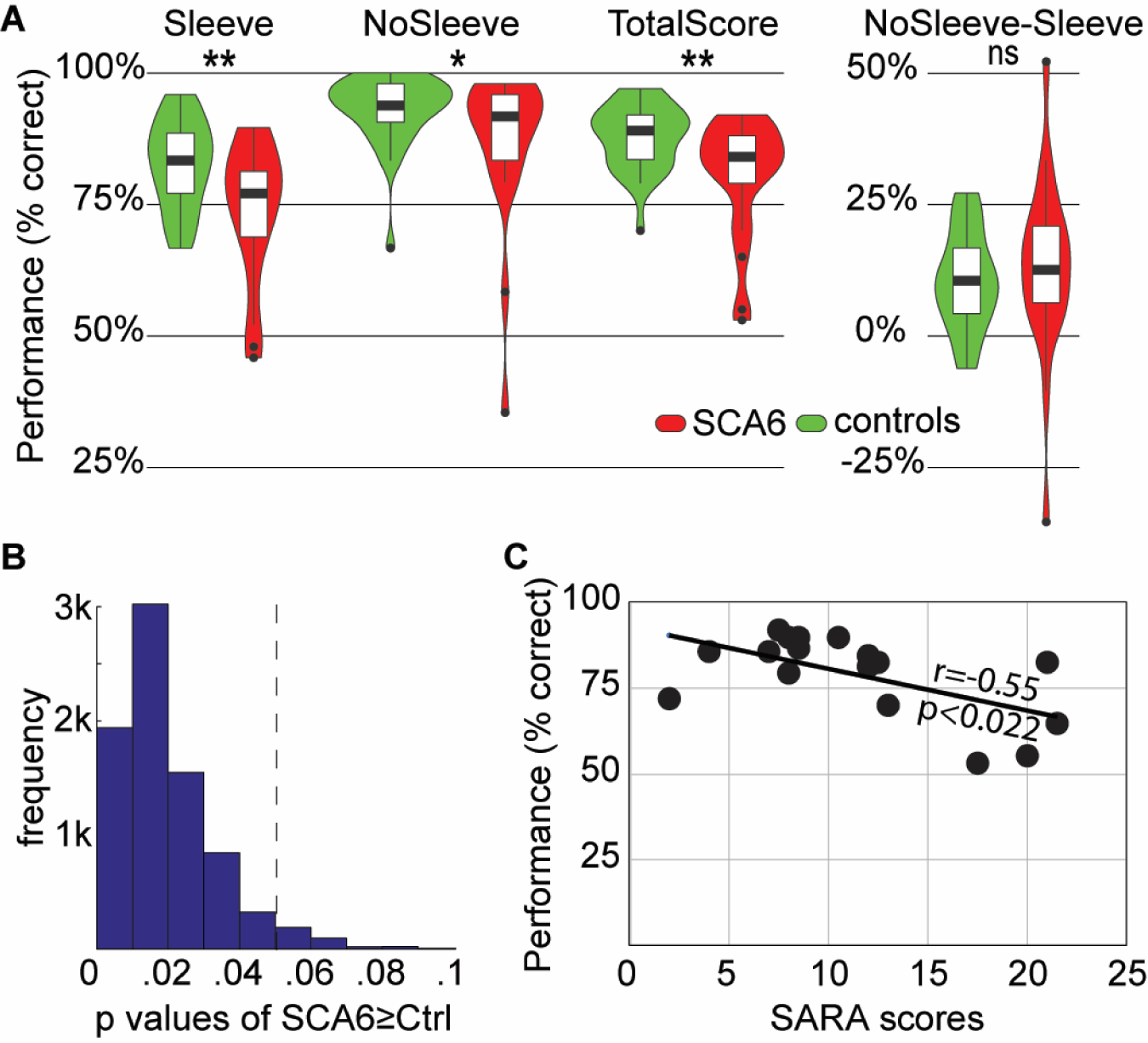
**(A)** Violin plot of the performance (percent correct responses) in the weight-discrimination task for the 21 SCA6 patients (red) and 31 controls (green) for the different conditions. *:p<0.05, **:p<0.01 using Mann-Witney U-tests to compare SCA6 vs controls group in each condition. **(B)** Distribution of p-values obtained from the 8008 possible subsamples of gender matched control groups, again using the Mann-Whitney U test to compare the total score (Sleeve and NoSleeve trials together) across groups. **(C)** The significant negative association between symptom severity (SARA) and total score in the weight perception task. The r-value reflects the non-parametric Spearman rank-order correlation. Higher SARA scores reflect more severe symptoms and predict more perceptual impairment.

To explore if this group difference in the performance could be due to the less than ideal matching on gender, we performed two further analyses. First, we performed a parametric ANOVA on the performance in the Sleeve condition with 2 Groups (SCA6 vs Ctrl) x 2 Genders. The interaction of Gender x Group was not significant (F_(1,48)_=2.66, p=0.11), suggesting that the group difference does not depend on gender. Second, we created control groups that were exactly matched in gender to the SCA6 group by sub-selecting 6 males out of the 16 available in the control group, keeping all the 15 females. There are 8008 ways to subsample 6 males out of 16, and for each of them, we calculated the p-value for the group difference in total performance using the Mann-Whitney U one tailed test. The median p-value across the 8008 subsamples was p=0.016 and 7675 of the 8008 (96%) had p<0.05 (Fig. 4B). This confirms that compared to the majority of randomly subsampled, gender matched control groups, the SCA6 group shows impaired performance in our task.

To explore whether there is a significant association between the severity of the degenerative disorder and the performance in our task, we calculated the Spearman rank order correlations between the total performance score and the SARA score for the 17 patients for which we do have the SARA score (Fig. 4C). We found that the association is significant: R=-0.55, t_(15)_=-2.54, p<0.022.

Finally, to explore whether the perceptual impairment we observe in SCA6 patients would also be visible in implicit measures, we added eye tracking in our last participants (4 SCA6 and 7 controls), which did not show any significant group difference (Supplementary results 2.3). Given the small sample size only large group differences could have been detected, however, the qualitatively similar pattern in the two groups suggests that SCA6 did not severely alter how subjects explored the stimuli in space and time.

## Discussion

Our primary aims were (a) to explore whether and where the cerebellum is robustly activated by the observation of other individual’s hand actions, of others, and (b) whether disrupting the cerebellum leads to significant impairments in hand action observation.

Regarding activations, using scanning parameters that include the entire cerebellum (both in terms of field of view during acquisition and bounding box during analysis) we found that across three studies and a total of 79 participants, the cerebellum was consistently recruited by the contrast between goal-directed hand actions and meaningless movements of the hand close to an object. Single subject analyses confirmed that the cerebellum was recruited in all but 4 participants. More specifically, we find that activity is reliably induced in the lateral hemispheres of lobule VI, and in a cluster including Lobules VIIb and VIIIa. All these activations are bilateral. Without using smoothing, it is apparent that the dorsal cluster in Lobule VI is distinct from activity in the ventral visual pathway, and is thus not the result of bleeding of activity from visual neocortical regions. Each of these clusters were found to be activated in the majority of individual participants. Together these results provide strong evidence that the cerebellum is consistently recruited by hand action observation.

This raises the question of why former studies failed to consistently report cerebellar activations. Our comparison of pipelines identifies two potential reasons: (i) up to SPM8, the default bounding box for analyses prevented the identification of some of the cerebellar clusters, and (ii) most studies focusing on the cerebrum have to choose between a larger field of view (i.e. more spatial coverage) vs. a shorter acquisition time (i.e. increased task sensitivity), which often ends in favouring a smaller field of view therefore cutting out the cerebellum in at least some participants. At the second level of analysis, if part of the cerebellum is missing in the field of view for some of the participants, this region is entirely removed from the search volume on which statistical analyses are computed across all subjects. This may have further reduced the consistency with which cerebellar activity is reported. Finally, a comparison between the number of participants activating our cerebellar ROIs compared to classic cerebral ROIs such as BA44 or PF, showed that the cerebellar ROIs indeed are slightly less reliably recruited, providing an additional factor. Overall, our three studies provide clear evidence that with proper measurement procedures and analyses pipelines, cerebellar recruitment during hand action observation can be demonstrated. The finding that these same regions are also activated when using a different, weight judgement task (Exp. #4) in our sample shows that this consistency does not depend on a specific task.

It is interesting that the specific locations of consistent activations in our study are overlapping with or adjacent to regions that have been associated with the sensorimotor control of hand actions in the cerebellum (Schlerf *et al*., 2015). One of our foci was localized in the anterior part of lobule VI, which is associated with the primary sensorimotor map of finger motions in the cerebellum (Grodd *et al*., 2001; Schlerf *et al*., 2015). Our second focus was localized in the posterior inferior lobule VIIb expanding into lobule VIIIa. Its location is spatially adjacent to the secondary sensorimotor finger map (Grodd *et al*., 2001; Schlerf *et al*., 2015). This is in line with the notion that cerebro-cerebellar loops involved in fine kinematic control of hand actions may also serve as a valuable system to process fine kinematics of observed actions (Miall, 2003; Wolpert, Doya and Kawato, 2003; Fuentes and Bastian, 2007; Gazzola and Keysers, 2009; Rizzolatti and Sinigaglia, 2010; Sokolov, Miall and Ivry, 2017).

To explore whether the cerebellum is necessary for extracting information from the kinematics of the hand actions of others, we tested whether patients with SCA6 are impaired in a weight-lifting task that has been shown to depend on precise processing of hand movement kinematics(de C. Hamilton *et al*., 2007). Our results indicate that SCA6 patients are indeed impaired in their kinematic processing as borne out by a group difference in the Sleeve condition that impoverishes muscle shape information. This impairment was more pronounced in patients with more severe SCA6 symptoms. Interestingly, when we analysed the data of the paradigms without the sleeves, we found that muscle shape processing appears to be preserved, as Bayesian statistics confirmed that the patients benefited from the additional muscle shape as much as the controls did. These results complement the results of the only other study that has, to our knowledge, examined the impact of cerebellar damage in action observation (Cattaneo *et al*., 2012) in that the two studies probed different aspects of hand action observation. In the task of Cattaneo, participant viewed four still frames of an action, and had to decide which was not part of that action. Solving that task does not require fine kinematic analyses, but an understanding of whether a particular hand-object interaction would be appropriate to achieve a particular goal. In our task, all movies show a hand successfully lifting an object, and performance thus depends on analysis of kinematics. That SCA6 patients were impaired in the Sleeve condition, in which kinematics was the primary cue, but could benefit from additional muscle shape, highlights that cerebellar degeneration particularly impairs kinematic processing. Moreover, these findings dovetail with our fMRI results, which show consistent cerebellar activity for the kinematic stimuli but not for the additional shape information provided in the NoSleeve condition.

As the cerebellum is involved in eye movement control, we were concerned that patients may be compromised in their ability to follow the movements of the arm with their gaze. However, our control data obtained from a small number of SCA6 patients does not suggest severe impairments in how our patients deploy their gaze. Future studies could include fMRI of SCA6 patients to explore where in the cerebellum degeneration alters task-related activity, and whether this includes regions associated with gaze-control. A previous VBM study points to a loss of gray matter in the hemispheres of lobule VI as the primary cause of upper limb ataxia triggered by SCA6 (Rentiya *et al*., 2017), which is in close vicinity to and partly overlaps with regions in which we find cerebellar activations to action observation, but is lateral relative to the sections of lobule VI mostly associated with eye movements (Supplementary Fig. 3).

Because interfering with one node of the action observation network is known to disrupt the activity in the other connected nodes (Valchev *et al*., 2016), this should not be taken as evidence that the cerebellar activity, per se, is necessary for the processing of observed actions. It could be that SCA6 disrupted activity in other connected brain regions that in turn are necessary for the conscious report of weight. Instead, our findings should be interpreted at the network level, to suggest that the cerebellum is a necessary node in a network that contributes to optimal perception and interpretation of observed hand actions.

In the light of our findings we believe that it is time to consider the cerebellum a reliable and necessary component of the network that allows us to process the kinematics of observed hand actions. Clinically, one of the core complaints of many stroke survivors and their spouses are impairments in social cognition (Hillis, 2014). These social sequelae are often not on the radar of neurological staff. We hope that by showing that SCA6 patients have deficits in perceiving the kinematics of the actions performed by other individuals – deficits that gets worse with the severity of the disease - our results contribute to an increased awareness of the social consequences of neurological disorders affecting the cerebellum. Being impaired in perceiving what other individuals around us do is likely to impact the way we related to others and as thereby reduce our wellbeing.

## Supporting information

## Acknowledgments

We thank Filippo Migliorati and Rajat Thomas for their support during data analysis. We thank Marc Thioux, Anita Sibeijn-Kuiper, Judith Streurman and Teresa De Sanctis for help in acquiring the MRI data. We thank Jessica Willems, Esther Brusse, Henk-Jan Boele, Ruben van der Giessen, Cullen B. Owens for facilitating the experiments with SCA6 patients and performing neurological examinations. We thank the Spinoza Center Amsterdam and the Neuroimaging Center at the UMCG for contributing scan time for the fMRI data acquisition. We finally thank F. Van Overwalle for providing the mirror-maps from his meta-analysis (Van Overwalle *et al*., 2014). **Author contribution.** Funding was obtained by V. Gazzola, C. Keysers, C. De Zeeuw and R. Bhandari. The experiment was conceived by V. Gazzola, C. De Zeeuw and C. Keysers together with A. Abdelgabar and J. Suttrup. FMRI data were collected by J. Suttrup (Exp.#1) and R. Bhandari (Exp.#2-4) with facilitation from V. Gazzola. Behaviuoral data on healthy volunteers and SCA6 patients were collected by A. Abedlgabar, R. Broersen and S. Picard with faciliation of C. De Zeeuw. Data analysis was performed by J. Suttrup, R. Bhandari, A. Abdelgabar, R. Broersen and V. Gazzola, with suggestions from C. Keysers and C. De Zeuuw. The manuscript was written by J. Suttrup, A. Abdelgabar, R. Broersen and V. Gazzola, with extensive input from C. Keysers and comments from all the other authors.

## Table legends

**Table 2. Cerebellar activations to ActionOBS-CtrlOBS for Exp. #1 to #3**. Regions with ActionOBS-CtrlOBS≥4.31 labelled using SPM Anatomy Toolbox. Results are shown at p_FWE_ < 0.05 with cluster size >10 voxels. From left to right: the cluster size in number of voxels, the number of voxels falling in a cyto-architectonic area, the percentage of the cluster that falls in the cyto-architectonic area, the hemisphere (L=left; R=right), the name of the cyto-architectonic area when available or the anatomical description, the percentage of the area that is activated by the cluster, the t values of the peaks associated with the cluster followed by their MNI coordinates in mm.

**Table 3: Comparison between Exp #1, #2 and #3 in number of voxel and peak distance per cluster of activity.** For each of the four cerebellar clusters, and for each experiment separately, the number of voxels surviving p_FWE_<0.05 for the contrast ActionOBS-CtrlOBS is reported. The fourth column reports the number of voxels counted within the conjunction of the three experiments. The last row indicates the number of cerebellar voxels not falling within the region of interest. Columns 5-7 indicate the minimum Euclidean distance between the activation-peaks identified belonging to the four clusters by the Anatomy toolbox for SPM.

**Table 4. Cerebellar activations to the weight discrimination task**. Results are shown at p_FWE_< 0.05 with cluster size >10 voxels. Conventions as in Table 2.

