## Supplementary material for "Action perception recruits the cerebellum and is impaired in spinocerebellar ataxia patients"

### Supplementary tables and figures

**Supplementary Table 1: Scanning parameters across fMRI experiments.** MB=multi band (i.e. number of simultaneously acquired slices), S=sense factor. TR=repetition time in msec. Exp.=experiment.

| | | **Groningen, NL, Exp. #1** | **SPINOZA, Amsterdam, NL Exp. #2 & #4** | | | | **SPINOZA, Amsterdam, NL Exp. #3** | | | | |
|---|---|---|---|---|---|---|---|---|---|---|---|
| **Anatomical acquisition parameters** | Sequence | 3D-spoiled T1-weighted | T1-weighted | | | | T1-weighted | | | | |
| | Slices | 170 | 170 | | | | 250 | | | | |
| | Resolution (mm) | 256x256 | 240x222 | | | | 256 x256 | | | | |
| | Field of view (mm) | 232 | 240 x 240 x 170 | | | | 240x256x250 | | | | |
| | Voxel size (mm) | 1x1x1 | 1x1x1 | | | | 1x1x1 | | | | |
| **Functional acquisition parameters** | Sequence | T2*-weighted | T2*-weighted | | | | T2*-weighted | | | | |
| | Slices | | MB1S2 Exp#2&4 | MB2S2 Exp#2 | MB3S2 Exp#2 | MB4S2 Exp#2 | MB1S2 | MB1S2 | MB2S2 | MB4S1.5 | MB4S2 |
| | Nº of slices | 41 | 40 | 40 | 39 | 40 | 44 | 36 | 44 | 44 | 44 |
| | Echo time (ms) | 28 | 27.6 | 27.6 | 27.6 | 27.6 | 30 | 30 | 30 | 30 | 30 |
| | Thickness (mm) | 3.5 | 3.5 | 3.5 | 3.5 | 3.5 | 2.7 | 3.3 | 2.7 | 2.7 | 2.7 |
| | Gap (mm) | NO | 0.34 | 0.34 | 0.34 | 0.34 | 0.27 | 0.33 | 0.27 | 0.27 | 0.27 |
| | Flip angle | 70 | 72.9 | 72.9 | 72.9 | 72.9 | 79 | 75 | 64 | 51 | 50 |
| | Repetition time | 2000 | 2060 | 1230 | 760 | 570 | 2450 | 2000 | 1220 | 700 | 630 |
| | Resolution | 64x62 | 80x157 | | | | 80x78 | | | | |
| | Field of view (mm) | 224 | 240 x 240 x 153.65 | | | | TR[2450,1220,700,630]=216x216x130.4<br>TR[2000]=240x240x130.3 | | | | |
| | Voxels size (mm) | 3.5x3.5x3.5 | 3.0x3.0x3.0 | | | | TR[2450,1220,700,630]=2.7x2.7xx2.7<br>TR[2000]=3.0x3.0x3.3 | | | | |
| | Number of volumes acquired | 345 | 360 Exp.#2; 325 and 326 for session 1 and 2 of Exp. #4 | | | | 200 | 245 | 400 | 700 | 780 |

**Supplementary Table 2: Schema of the temporal order of processing steps and bounding box size for the four processing pipelines testing the effect of different normalization parameters in Exp. #1.** Each column details the order of processing steps used in a specific pipeline. All pipelines start with core preprocessing including: centering of the anatomical image to the anterior commissure, slice timing, realignment of all EPI images to one another. Finally, the T1 image is coregisted to the mean EPI image. Only then do the different pipelines start to differ. The acronym of the different pipelines is constructed using the following elements. WB: whole-brain. Cereb: Cerebellum specific analysis. GLM: general linear model, referring to the first level of analysis. SUIT: spatially unbiased atlas template. MNI: MNI brain template. norm: normalization.

| WB_MNInorm_GLM | WBcut_MNInorm_GLM | Cereb_GLM_SUITnorm | WB_GLM_MNInorm |
|---|---|---|---|
| slice timing<br>EPI realignment<br>T1-EPI coregistration | slice timing<br>EPI realignment<br>T1-EPI coregistration | slice timing<br>EPI realignment<br>T1-EPI coregistration | slice timing<br>EPI realignment<br>T1-EPI coregistration |
| **whole-brain normalization**<br>**large** box | **whole-brain normalization**<br>**small** box | GLM | GLM |
| smoothing | smoothing | **cerebellar normalization**<br>**large** box | **whole-brain normalization**<br>**large** box |
| GLM | GLM | smoothing | smoothing |

**Supplementary Table 3. Cerebellar activations to ActionOBS-CtrlOBS for the WB_MNInorm_GLM and the Cereb_GLM_SUITnorm pipelines.** Regions with ActionOBS-CtrlOBS≥5.4. Only clusters with minimum 10 voxels are reported. Clusters are described using SPM Anatomy Toolbox. From left to right: the cluster size in number of voxels, the number of voxels falling in a cyto-architectonic area, the percentage of the cluster that falls in the cyto-architectonic area, the hemisphere (L=left; R=right), the name of the cyto-architectonic area when available or the anatomical description, the percentage of the area that is activated by the cluster, the t values of the peaks associated with the cluster followed by their MNI coordinates in mm.

| Cluster size | # Voxels in cyto | % Cluster | Hem | Cyto or anatomical description | % Area | Peak Information | | | |
|---|---|---|---|---|---|---|---|---|---|
| | | | | | | T | x | y | z |
| **Exp. #1,WB_MNInorm_GLM pipeline, ActionOBS-CtrlOBS $p_{FWE}$<0.05, t=5.8, min. 10 voxels** | | | | | | | | | |
| 138 | 130.8 | 94.7 | R | Lobule VI (Hem) | 7.2 | 9.41 | 30 | -56 | -24 |
| | | | | | | 8.13 | 32 | -50 | -30 |
| | | | | | | 6.96 | 38 | -48 | -28 |
| | | | | | | 6.95 | 30 | -46 | -24 |
| 100 | 66.6 | 66.6 | L | Lobule VIIIa (Hem) | 8.8 | 7.48 | -26 | -60 | -52 |
| | | | | | | 6.41 | -20 | -64 | -56 |
| | 27.8 | 27.8 | L | Lobule VIIb (Hem) | 4.1 | 7.68 | -12 | -74 | -48 |
| | 4.5 | 4.5 | L | Lobule VIIIb (Hem) | 0.7 | | | | |
| 94 | 83 | 88.3 | R | Lobule VIIb (Hem) | 12.7 | 9.28 | 18 | -78 | -52 |
| | | | | | | 9.13 | 14 | -76 | -48 |
| | | | | | | 8.63 | 16 | -74 | -46 |
| 79 | 77.5 | 98.1 | L | Lobule VI (Hem) | 4.1 | 12.22 | -28 | -54 | -24 |
| | | | | | | 6.24 | -34 | -52 | -28 |
| 47 | 43.1 | 91.8 | R | Lobule VI (Hem) | 2.4 | 10.13 | 22 | -72 | -22 |
| 47 | 45.8 | 97.3 | R | Lobule VIIIa (Hem) | 6.3 | 7.84 | 24 | -64 | -54 |
| | | | | | | 7.03 | 30 | -60 | -52 |
| | | | | | | 6.5 | 18 | -68 | -54 |
| **Exp. #1, Cereb_GLM_SUITnorm pipeline, ActionOBS-CtrlOBS $p_{FWE}$<0.05, t=5.4, min. 10 voxels** | | | | | | | | | |
| 214 | 199.1 | 93 | R | Lobule VI (Hem) | 11 | 9.18 | 30 | -56 | -24 |
| | | | | | | 8.25 | 22 | -72 | -22 |
| | | | | | | 7.48 | 32 | -52 | -28 |
| | | | | | | 6.72 | 26 | -66 | -20 |
| 159 | 110.4 | 69.4 | L | Lobule VIIIa (Hem) | 14.5 | 7.97 | -26 | -58 | -54 |
| | | | | | | 7.62 | -24 | -60 | -52 |
| | | | | | | 7.11 | -20 | -62 | -58 |
| | 25.4 | 16 | L | Lobule VIIb (Hem) | 3.7 | 8.35 | -16 | -74 | -52 |
| | | | | | | 8.05 | -14 | -72 | -48 |
| | 21.1 | 13.3 | L | Lobule VIIIb (Hem) | 3.5 | 8.44 | -20 | -60 | -48 |
| 118 | 97.4 | 82.5 | R | Lobule VIIb (Hem) | 14.9 | 8.35 | 16 | -74 | -46 |
| | | | | | | 7.93 | 14 | -76 | -52 |
| | | | | | | 7.78 | 12 | -76 | -48 |
| | | | | | | 7.68 | 18 | -78 | -54 |
| | 14.1 | 12 | R | Lobule VIIIa (Hem) | 1.9 | | | | |
| 101 | 99 | 98 | L | Lobule VI (Hem) | 5.3 | 11.04 | -28 | -54 | -24 |
| | | | | | | 7.35 | -24 | -56 | -28 |
| | | | | | | 6.08 | -28 | -60 | -22 |
| 72 | 65.6 | 91.1 | R | Lobule VIIIa (Hem) | 9 | 8.17 | 24 | -64 | -54 |
| | | | | | | 7.56 | 26 | -60 | -56 |
| **Exp. #1, WB_GLM_MNInorm pipeline, ActionOBS-CtrlOBS $p_{FWE}$<0.05, t=5.4, min. 10 voxels** | | | | | | | | | |
| 102 | 91.4 | 89.6 | | Lobule VIIb (Hem) | 14 | 9.3 | 18 | -78 | -52 |
| | | | | | | 8.97 | 14 | -76 | -48 |

| | | | | | | | | |
|---|---|---|---|---|---|---|---|---|
| | | | | | 8.52 | 16 | -74 | -46 |
| 101 | 96.3 | 95.3 | Lobule VI (Hem) | 5.3 | 9.3 | 30 | -56 | -24 |
| | | | | | 8.06 | 32 | -50 | -30 |
| | | | | | 7.1 | 36 | -50 | -30 |
| | | | | | 6.91 | 30 | -46 | -24 |
| 94 | 61.6 | 65.6 | Lobule VIIIa (Hem) | 8.1 | 7.52 | -26 | -60 | -52 |
| | | | | 4 | 6.4 | -26 | -64 | -56 |
| | | | | 0.7 | 6.16 | -22 | -64 | -56 |
| | 27 | 28.7 | Lobule VIIb (Hem) | | 7.74 | -14 | -74 | -48 |
| 73 | 71.5 | 97.9 | Lobule VI (Hem) | 3.8 | 12.49 | -28 | -54 | -24 |
| 42 | 41.5 | 98.8 | Lobule VIIIa (Hem) | 5.7 | 7.74 | 24 | -64 | -54 |
| | | | | | 7.02 | 30 | -60 | -52 |
| | | | | | 6.54 | 18 | -68 | -54 |
| 39 | 35.9 | 92 | Lobule VI (Hem) | 2 | 10.13 | 22 | -72 | -22 |
| | | | | | 7 | 26 | -66 | -20 |

**Supplementary Table 4. Number of activated cerebellar voxel.** Median of activated voxels for the ActionOBS-ActionCtrl contrast for each experiment individually (Exp1-3) and when considered as a group (All; N=79), for each of the four anatomically defined cerebellar clusters identified at the group level (but separately for each hemisphere: L, left; R, right). Results are also reported for the whole cerebellum (Cereb), and for cortical areas activated by the same contrast and of similar volume (PF L, PF R, BA44L, BA44R). Percentage of participants having no voxels (% zeros) in any of the region of interest, and the percentage of those having at more than 10 voxels (%>10) are also reported. The size, in number of voxels, of each ROI is indicated under the ROI name in the first line.

| | | **VI L**<br>2085 | **VI R**<br>2041 | **VIIIa/ VIIb L**<br>1640 | **VIIIa/ VII R**<br>1588 | **Cereb**<br>21127 | **PF L**<br>2205 | **PF R**<br>2337 | **BA44 L**<br>869 | **BA44 R**<br>607 | **BA1/2 L**<br>1101 | **BA1/2 R**<br>1349 |
|---|---|---|---|---|---|---|---|---|---|---|---|---|
| **median** | **Exp1** | 14 | 47 | 4 | 5 | 208 | 517 | 373 | 115 | 41 | 115 | 501 |
| | **Exp2** | 4 | 10 | 9 | 54 | 186 | 244 | 146 | 89 | 20 | 88 | 254 |
| | **Exp3** | 67 | 50 | 8 | 82 | 384 | 450 | 282 | 91 | 72 | 91 | 705 |
| | **All** | 14 | 31 | 7 | 44 | 224 | 379 | 282 | 92 | 30 | 92 | 485 |
| **% zeros** | **Exp1** | 25.8 | 25.8 | 45.2 | 29.0 | 12.9 | 0.00 | 0.00 | 12.9 | 19.4 | 12.90 | 0.00 |
| | **Exp2** | 32.0 | 24.0 | 28.0 | 12.0 | 0.0 | 0.00 | 8.00 | 4.0 | 20.0 | 4.00 | 4.00 |
| | **Exp3** | 17.4 | 21.7 | 26.1 | 17.4 | 0.0 | 8.70 | 13.04 | 13.0 | 13.0 | 13.04 | 4.35 |
| | **All** | 25.3 | 24.1 | 34.2 | 20.3 | 5.1 | 2.53 | 6.33 | 10.1 | 17.7 | 10.13 | 2.53 |
| **% >10** | **Exp1** | 51.6 | 58.1 | 38.7 | 45.2 | 77.4 | 100.0 | 93.55 | 83.9 | 67.7 | 83.87 | 100.00 |
| | **Exp2** | 40.0 | 48.0 | 48.0 | 84.0 | 88.0 | 96.00 | 84.00 | 92.0 | 60.0 | 92.00 | 92.00 |
| | **Exp3** | 60.9 | 69.6 | 47.8 | 73.9 | 95.7 | 91.30 | 82.61 | 73.9 | 78.3 | 73.91 | 91.30 |
| | **All** | 50.6 | 58.2 | 44.3 | 65.8 | 86.1 | 96.20 | 87.34 | 83.5 | 68.4 | 83.54 | 94.94 |

**Supplementary Fig. 1: Effect of different analysis pipelines on cerebellar activation to action observation. (A-D)** Cerebellar activations ($p_{FWE}<0.05$) for the four pipelines displayed on flat maps of the cerebellum. Color code in (A) identifies the different cerebellar lobules (Diedrichsen and Zotow, 2015). **(E)** For each of the four pipelines (A-D), the graphs shows the mean top 5% of t-values, and indicates the significant differences between the traditional pipeline (A) and the other three (B-D). *p<0.002, **p<0.001, ns=not significant. **(F)** As in (A) on the uncolored flat-map and on a sagittal slice. When smoothing is applied, the cerebellar cluster marked with the black arrow is part of a temporal lobe cluster. When smoothing is not applied as in (G) the cluster is clearly cerebellar. **(G)** Same as in (A) and (F) but computed on unsmoothed data.

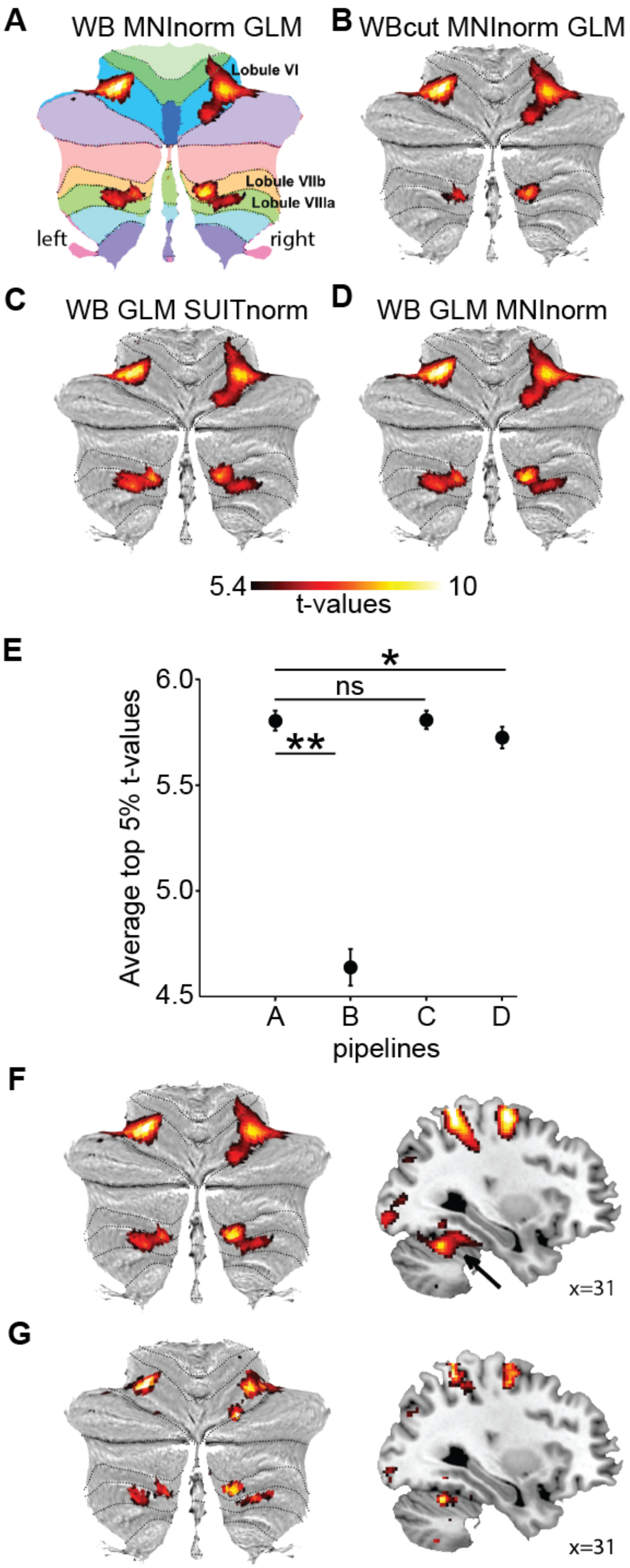

**Supplementary Fig. 2**: **Task-related eye movement dynamics. (A, B)** Heatplots based on spatial gaze intensity, identifying IRs during arm movement in the task for each group per condition. Both controls and SCA6 subjects focused significantly on the proximal arm muscles in the NoSleeve condition, although they focused on both distal and proximal part of the arm in the Sleeve condition, with an IR around the wrist. **(C)** Gaze position along the vertical axis throughout the task, showing tracking of upward going arm movements for both groups during both conditions. **(D)** Gaze position along the horizontal axis shows proximal arm focusing during arm movements in the NoSleeve condition. **(E, F)** Total gaze distance in the Sleeve condition along the V and H axis, respectively, shows no group differences. **(G, H)** Total gaze distance in the Nosleeve condition along the V and H axis, respectively, also shows no difference between groups.

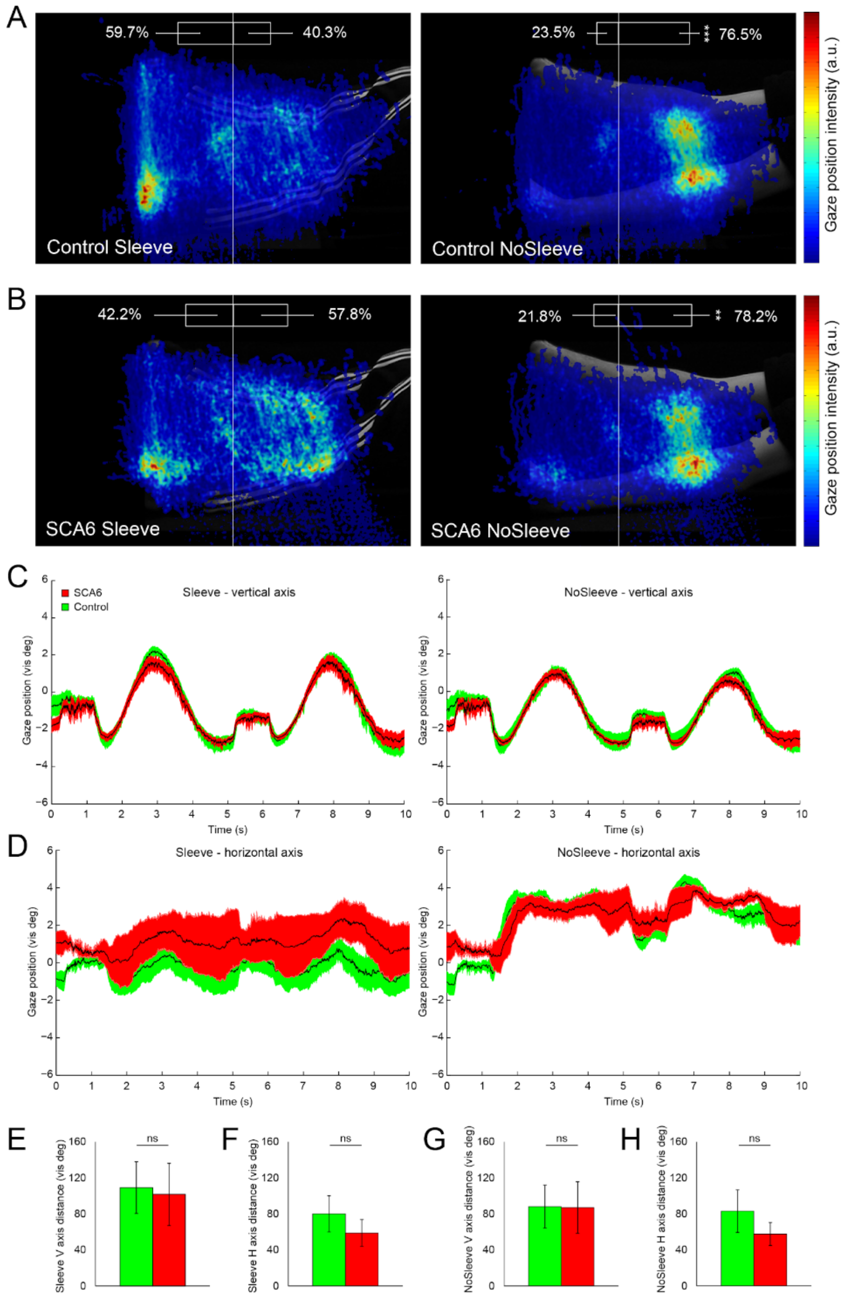

**Supplementary Fig. 3**: **Comparison with locations of VBM changes and eye movement cerebellar activity**. Red: location of cortical and cerebellar activity during eye movement tasks as identified by a Neurosynth (http://neurosynth.org/) meta-analysis with the term 'eye movements' (116 studies and 5486 activation clusters identified). Blue: cerebellar activation maps of the global null conjunction at pFWE<0.05 of the Sleeve and NoSleeve conditions falling within the conjunction of the ActionOBS-ActionCtrl contrasts of Exp.#1,2 and 3. From top to bottom, the coordinates (and blue crosses when present) indicate the location of: the left (top row) and right (middle row) lobule VI peak of correlation between VBM and SCA6 patients' performance in the Grooved Pegboard (from Table 3 in (Rentiya et al., 2017)), and (bottom row) the lobule VI peak of activity to eye movements as identified by the met-analysis in Neurosynth. Note how the VBM results (crosses) associated with SCA6 overlap with the regions recruited by our task (blue) but fall lateral to the regions most involved in eye movements (red). L: left hemisphere. R: right hemisphere. A: anterior. P: posterior. The clusters of activity are shown on the ch2better template from MRIcron (https://www.nitrc.org/projects/mricron/).

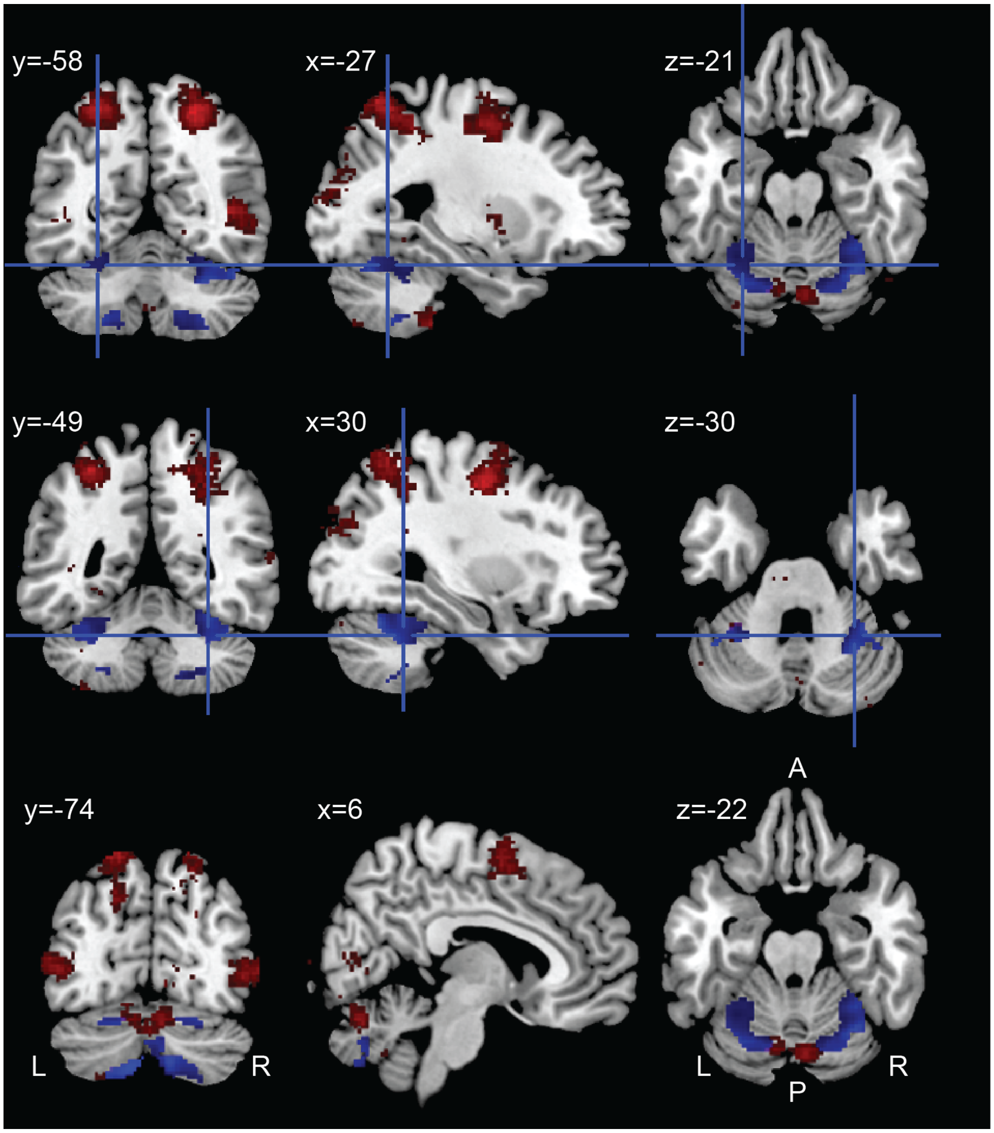

### Supplementary materials and methods

*Supplementary method 1.0. Recruitment procedure*

The SCA6 patient group was recruited in collaboration with the department of Neurology at the Erasmus University Medical Center Rotterdam. A number of SCA patients had been approached by their physician for participating in a previous research project. As such, we built a database with names and addresses of volunteers that were willing to participate in research. Several volunteers approached our contact person individually and they were added to the database. Moreover, several participants were member of the Dutch society for Autosomal Dominant Cerebellar Ataxia (ADCA; www.ataxie.nl). Twenty-nine healthy control participants were recruited as relatives or friends of SCA patients or investigators, and were selected based on their age and gender to match the SCA patient group. All participants received an information letter or email asking whether they were interested in participating in the study.

*Supplementary method 1.1. Impact of different analysis pipelines*

Four pipelines were computed and compared to test the effect of different spatial normalization procedures. The analyses were done in SPM8 (Wellcome Trust Centre for Neuroimaging, UCL, UK) and complemented with customized Matlab scripts (Matlab 7.14; The MathWorks Inc., Natick, USA). Common to all, data pre-processing included: slice time correction of functional images using the bottom slice, located in the posterior cerebellum, as a reference slice; realignment and co-registration of the T1-weighted anatomical to the mean functional image. Below a description of the steps that followed for each pipeline separately. Supplementary Table 2 more schematically illustrates the analysis steps used for the four pipelines. For Exp. #1, the acquisition plane was tilted by 30-45° from the AC-PC plane to cover the entire cerebellum.

Pipeline I: WB_MNInorm_GLM. As commonly done in fMRI analyses, the whole brain (WB) functional images were brought to MNI space before computing the GLM, using the normalization (norm) parameter generated during segmentation of the anatomical image (final voxel size: 2 × 2 × 2 mm). We manually adjusted the SPM8 bounding box settings to [-90 -126 -72; 91 91 109] to cover the entire cerebellum.

Pipeline II: WBcut_MNInorm_GLM. Same as for Pipeline I but without adjustment of the bounding box, which was left to the default SPM8 settings [-78 -112 -50; 78 76 85]. This allowed us to identify which part of the activation was left out in previous studies focusing on the cortex.

Pipeline III: Cereb_GLM_SUITnorm. A template for cerebellar-specific normalization using a high-resolution atlas of the human cerebellum (Cereb) is available in the literature (SUIT, Diedrichsen, 2006; Diedrichsen *et al.*, 2009), and it has been shown to improve the alignment of anatomical landmarks and increase average t-values for cerebellar functional data sets (Diedrichsen, 2006). The SUIT template is in MNI space, but it is based on a group of 20 participants in order to create an average anatomical template with enough anatomical details to account for the small size of cerebellar functional regions(Diedrichsen, 2006). In pipeline III, following the method suggested by (Diedrichsen, 2006), after co-registration, the functional images were directly fed into the subject-level general linear models. Resulting contrast images were subsequently normalized to the SUIT-space, by first isolating the cerebellum from the T1 images using an automated algorithm from the SUIT toolbox (www.icn.ucl.ac.uk/motorcontrol/imaging/suit.html). This step resulted in a cropped anatomical image covering the cerebellum and adjacent cortical regions, which was normalized into SUIT cerebellar space. The transformation parameters obtained during the normalization were then used to normalize the contrast images resulting from the first level GLM (final voxel size: 2 × 2 × 2 mm).

Pipeline IV: WB_GLM_MNInorm. The Cereb_GLM_SUITnorm pipeline differs from traditional whole brain pipelines not only in the normalization template but also in the moment at which normalization is computed: after vs. prior to the GLM. In order to assess the impact of this difference, pipeline IV was run with the same temporal order used in pipeline III: the first level GLM used the co-registered functional images, and the normalization to the MNI whole brain template was applied on the contrast images resulting from the GLM. Normalization parameters were calculated during the segmentation of the whole-brain T1 anatomical image (final voxel size: 2 × 2 × 2 mm). In order to include the entire cerebellum, the bounding box was manually adjusted to [-90 -126 -72; 91 91 109].

Smoothing, using a 6 mm FMHW Gaussian kernel, was applied to each pipeline after normalization. Although spatial smoothing is routinely applied in the neuroimaging literature, it poses the possibility of leakage of activation between the anterior cerebellum and the temporal cortex. We therefore report our results with and without the 6 mm FMHW Gaussian filter for our WB_MNInorm_GLM pipeline, which is closest to traditional MRI analysis in the literature. The impact of smoothing on leakage between cortical and cerebellar activation is also investigated by comparing cerebellar activity resulting from whole brain analyses run on either smoothed or unsmoothed data.

The same general linear models was applied to each pipeline. Two standard box car predictors modelled the ActionOBS, CtrlOBS and static conditions, and were convolved with the canonical hemodynamic response function (HRF). The last six regressors of no interest included the displacements and rotations determined during image realignment. The ActionOBS-CtrlOBS contrast was computed at the subject-level to generate action specific activations for observation, and tested against zero with a one-sample t-test at the group level. The static condition was modelled at the first level, but for the purpose of this study, not analyzed at the second level.

The choice of the statistical threshold at which to report the group results is not trivial. First, we want to be able to compare multiple activation maps resulting from different preprocessing pipelines. Second, correction algorithms based on random field theory require a certain amount of smoothness (Brett et al. 2004), which is not given using unsmoothed data sets. All statistical maps thus are thresholded at $p_{FWE}<0.05$ and have minimal cluster size of 10 voxels. We chose peak-level FWE-correction because we wished to (i) interpret activation of individual voxels, and, motivated by the inconsistencies of cerebellar activations in the literature, (ii) limit the risks of Type I errors.

Anatomical descriptions of cerebral activity were guided by the probabilistic cytoarchitectonic maps (Geyer *et al.*, 1996, 2000; Amunts *et al.*, 1999; Geyer, Schleicher and Zilles, 1999; Grefkes *et al.*, 2001; Geyer, 2004; Eickhoff *et al.*, 2005, 2006; Caspers *et al.*, 2006; Choi *et al.*, 2006) implemented in the anatomy toolbox for SPM (http://www.fz-juelich.de/ime/spm_anatomy_toolbox) (Eickhoff *et al.*, 2005, 2006, 2007).

To investigate the impact of different pipelines, we used unsmoothed contrast images resulting from the 1$^{st}$-level analysis of action observation data after setting them to identical image dimensions (91 × 109 × 91) using the ImCalc function. The contrast images were then fed to group-level, one-sample t-test, GLM models (ActionOBS-CtrlOBS>0), one for each pipeline. Voxels missing in any of the four pipelines were excluded for the analysis with exception of the voxels missing due to the bounding box size, which were coded with 0, such that differences between the pipelines could also be evaluated in the inferior posterior cerebellum. The t-values from the four group-level t-values cerebellar maps were summed up using the NIfTI toolbox (version 1.25). In line with (Diedrichsen, 2006), in order to compare the results of the four pipelines without biasing the comparison a priori to any specific pipeline, we selected from the sum of the four t-maps, the location of the 5 percent of voxels with the highest t-value sums as voxels of interest. The pipelines were then compared using a repeated-measures ANOVA design that considers each voxel of interest as a 'subject', and each pipeline a repeated measurement of this 'subject' (i.e. voxel). We then planned to perform t-tests that compare each of the pipelines against the WB_MNInorm_GLM pipeline, because the whole brain analysis is most frequently used in the neuroimaging literature.

*Supplementary method 1.2. Consistency maps*

To generate the consistency maps, the normalized, smoothed single-subject t-maps of action observation (ActionOBS-CtrlOBS>0) from Exp.#1-3 were thresholded at the t-value corresponding to $p_{unc}<0.001$ (T=3.1), which binarizes the images. All single-subject maps were then added together to generate the group-level consistency map, showing for each voxel the number of participants for which the voxel was significantly activated by action observation. The number of participants needed to show that a voxel is activated more consistently than expected by chance, was calculated using a cumulative binomial distribution with 31 repetitions and an associated probability of 0.001. The resulting probability was Bonferroni corrected using the number of voxels in the search volume (170675 for the whole brain). Thus, a voxel activated by four or more participants can be considered above chance (Gazzola and Keysers, 2009).

*Supplementary method 1.3. Eye-tracking data acquisition and analysis*

Eye tracking measures of 4 patients (mean age 60.4y ± 10.6 SD; mean SARA score: 10.88 ± 8.37 SD) and 7 control participants (mean age 63.5y ± 5.7 SD)  were collectedduring the weight discrimination task using the EyeLink® 1000 system (SR Research Ltd., Mississauga, Ontario, Canada) at 500Hz. Participants sat up straight behind a desk and kept their head placed on a Head Support (SR Research Ltd., Mississauga, Ontario, Canada), providing support for the chin and forehead during the entire task. Task stimuli were presented using PsychoPy2 (v1.84, UON, UK)(Peirce, 2009) on a 19 inch TFT monitor (UltraSharp 1907FP, Dell, TX, USA) with 300x380cm dimensions, a resolution of 1280x1080 pixels and at a refresh rate of 55Hz. Distance between participants' eyes and screen was 593 ± 22.8 mm and room background light was minimized during the task. The presented clips appeared in the middle of the presentation screen and covered 720x480 pixels. A 9-point calibration and validation was executed before starting the experiment. PsychoPy2 sent text messages to be registered in EyeLink at the beginning and end of each pair of clips for synchronization purposes.

EyeLink EDF files were converted into MATLAB-compatible (MathWorks, USA) variables using the 'edf2mat' script (JN van der Geest, Dept of Neurosci, Erasmus MC, Rotterdam) and were further analysed using custom-written MATLAB code. Data was obtained on eye-related events (i.e. blinks, fixations and saccades) by using default gaze parser settings (EyeLink 1000 User's

Manual, SR Research, Ontario, Canada). Data was filtered using a Gaussian lowpass filter with a 50Hz cutoff frequency and was converted from pixels to visual degrees, using X and Y resolution values as calculated by EyeLink. We synchronized eye movement recordings with start of each pair of clips based on message event timestamps corresponding to start of the pair of clips. Then, for each clip we visually determined periods in which object lifting-associated arm movement occurred and we quantified eye-related parameters of events occurring during those periods, comparing SCA6 patients and controls. Parameters included number of saccades, blinks and fixations, duration of blinks and fixations (in milliseconds), saccade peak velocity (in visual degrees/second), saccade amplitude (in visual degrees) and distance in the horizontal and vertical plain (calculated on trial basis during 'fixation periods', representing smooth pursuit and drift). All of these parameters were calculated based on clip periods where arm movement occurred. To make a distinction between distal and proximal part of the arm, we decided on a x-coordinate (pixel 380 from left side movie) based on movies where a arm without sleeve was visible. We took into account arm movement dynamics during the movie and attempted to encompass deformations of the brachior radialis muscle during lifting in the proximal part of the arm (right side of the movie) and movements of the hand and wrist in the distal part of the arm (left side). The same x-coordinate was used for every movie analysed in both Sleeve and NoSleeve conditions.

Heat plots (Supplementary Fig. 2A, B) were generated based on matrices where values represented summation of gaze positions for each datapoint *during the period of arm movement*, separating Sleeve versus NoSleeve conditions. Matrices were processed using a 2-D circular averaging filter with a radius of 3 in replicate boundary setting, converted into a grayscale image and thresholded using an alphamask, so that values higher than 0.002 (on a scale of 0-1) were plotted with 75% opacity using colormap 'jet' on top of a two merged representative frames from movies in the Sleeve and NoSleeve conditions, showing begin and end-position of a lift movement. Gaze position over time was calculated from gaze positions in the V and H axis of all subjects per group (Supplementary Fig 2C, D). Distance along both axis was calculated by summation of the absolute difference between sampling points during the task (Supplementary Fig. 2E - H). Figures were further processed in Illustrator CS6 (Adobe, USA).

The different eye tracking measures of patients and controls have been compared using pairwise independent two sample t-tests. While not being a prominent feature, previous studies have found abnormalities in saccades for SCA6 patients (Gomez *et al.*, 1997; Buttner *et al.*, 1998; Christova, Anderson and Gomez, 2008), based on this a priori hypothesis we have performed one-tailed tests for all saccade metrics. We have used FDR correction as a more lenient multiple comparisons correction method as opposed to the more conservative Bonferroni correction, to increase our sensitivity to group differences.

Eye tracking measurements have been performed on a subset of patients and controls, to check whether these subsets are representative for their respective groups we performed the main Group analysis on task performance. There were no significant performance differences between controls and their eye tracker subgroup [$F(1,36) = 0.6655$, $p > =0.420$, $w2 = 0.0038$] nor between patients and the eye tracker patient subgroup [$F(1,22) = 1.3605$, $p =0.256$, $w2 = 0.0097$].

### Supplementary Results

*Supplementary results 1.1 Effect of different analysis pipelines on cerebellar activation during action observation*

When we mapped the activations triggered by viewing goal directed hand actions compared to control stimuli (ActionOBS-CtrlOBS) with a traditional pipeline and a bounding box encompassing the whole cerebellum, we found four main clusters of activation. Supplementary Fig. 1A and Supplementary Table 3 locate these clusters in the bilateral Lobule VI, VIIIa and VIIb. When the smaller SPM8 default bounding box was used the activations in Lobule VIIIa were not visible as they were not included in the search volume used in the analyses (Supplementary Fig. 1B).

Analyzing the data with the specific cerebellar normalization and the procedure proposed by (Diedrichsen, 2006; Diedrichsen *et al.*, 2009) results in the same clusters of activity identified with the traditional approach (Supplementary Fig. 1C and Supplementary Table 3). When the average top 5% of t-values is compared between the traditional and cerebellar specific pipeline (Supplementary Fig. 1E), no significant difference is observed between the two approaches ($p>0.95$, $t=-0.06$). Bayesian paired sample t-test confirms that there is evidence for the two pipelines to give equal results ($BF_{10}=0.043$; https://jasp-stats.org/). To investigate the impact of running the GLM in the subject space, instead of on normalized data, as it is done in the cerebellar optimized pipeline, we re-calculated the whole brain analysis following the same order of pre-processing. While at visual inspection the maps look very similar (Supplementary Fig. 1D), the top 5% t-values is significantly lower ($p<0.002$, $t=3.14$). Supplementary Fig. 1E also indicates a significant drop of t-values ($p<0.001$, $t=11.8$) when the small bounding box is used, likely due to part of the active voxels not included in the statistical computation.

In summary, these results indicate that as long as the whole cerebellum is included in the analyses, activations are preserved across different analysis pipelines. Additionally in our data set, no clear advantage is observed when using the pipeline optimized for the cerebellum compared to the traditional one, possibly due to improvement of co-registration and normalization algorithms in the newer SPM releases, which do not make the specific adjustments for the cerebellum anymore necessary.

*Supplementary results 1.2 Effect of spatial smoothing on cerebellar activation during action observation*

Because the dorsal cerebellum is located close to the ventral temporal lobe, one concern in reporting cerebellar activations from whole brain analyses is that smoothing could make activations of the ventral visual stream bleed into the cerebellum. Comparing results computed on unsmoothed and smoothed data indicates that spatial smoothing increases the number of super-threshold voxels during action observation (ActionOBS-CtrlOBS>0) in the cerebellum by 168%, from 350 to 939 voxels, given the same t-value threshold of $t=5.8$ (corresponding to the most stringent t value resulting from FWE of whole brain smoothed and unsmoothed results). However, smoothing caused clusters that are separated when using unsmoothed data to merge into a single cluster (arrow in Supplementary Fig. 1F). In particular, the merging happened within the right cerebellar lobule VI, and bilaterally between the cerebellar lobule VIIb and VIIIa. Results on the unsmoothed data confirm cerebellar activations in all previously identified clusters, including the dorsal lobule VI, supporting the notion that activations are not the result of smoothing leading to a bleeding of activation from ventral visual cortex onto the adjacent cerebellum (Supplementary

Fig. 1G). Despite unsmooth results confirming the extensive cerebellar activation on the lobule VI, the cluster still belong to a bigger cluster encompassing the fusiform gyrus, making a clear attribution of voxels at the border to the fusiform or the cerebellum more difficult, which is evident in some of the tables. Additionally, 80% of the lobut VI cluster reported by (Van Overwalle *et al.*, 2014) falls within the fusiform regions (FG4 in particular), suggesting that smoothing might have had a bigger impact on the meta-analysis maps computation.

*Supplementary results 1.3 Eye movements during the weight discrimination task*

In the Sleeve condition, subjects from both groups focussed equally on the distal and proximal part of the arm (Ctrl: $t_{(12)}$=1.523, p=0.154; SCA6: $t_{(6)}$=-0.802, p=0.453; Supplementary Fig. 2A and B, left panels). In the NoSleeve condition both groups focussed significantly more on the proximal muscles of the lower arm (Ctrl: $t_{(12)}$=-9.482, p<0.001; SCA6: $t_{(6)}$=-4.238, p=0.005; Supplementary Fig. 2A and B, right panels). There was no group difference in either condition (Sleeve: $t_{(9)}$=1.112, p=0.295; NoSleeve: $t_{(9)}$=-0.197, p=0.848).

There were no significant group differences in any of the following parameters: number of saccades ($t_7$= 2.17, p-value= 0.197), Saccade peak velocity ($t_7$= 2.32, p-value= 0.197), saccade amplitude ($t_6$= 1.45, p-value= 0.327), duration of blinks ($t_5$= -0.15, p-value= 0.995), time lost blinking ($t_9$= 0.33, p-value= 0.995), duration of fixations ($t_6$= 1.72, p-value= 0.327), duration of fixation ($t_8$= -1.6, p-value= 0.327).

We next investigated whether the trajectory of eye movement over time differed across groups. (Supplementary Fig. 2C). We did not observe any differences between groups for the Sleeve (V: $t_{(9)}$=0.163, p=0.874; H: $t_{(9)}$=0.727, p=0.486; Supplementary Fig. 2E and F) and NoSleeve condition(V: $t_{(9)}$=0.03, p=0.977; H: $t_{(9)}$=0.762, p=0.465; Supplementary Fig. 2G and H).
